# AERON: Transcript quantification and gene-fusion detection using long reads

**DOI:** 10.1101/2020.01.27.921338

**Authors:** Mikko Rautiainen, Dilip A Durai, Ying Chen, Lixia Xin, Hwee Meng Low, Jonathan Göke, Tobias Marschall, Marcel H. Schulz

## Abstract

Single-molecule sequencing technologies have the potential to improve measurement and analysis of long RNA molecules expressed in cells. However, analysis of error-prone long RNA reads is a current challenge. We present AERON for the estimation of transcript expression and prediction of gene-fusion events. AERON uses an efficient read-to-graph alignment algorithm to obtain accurate estimates for noisy reads. We demonstrate AERON to yield accurate expression estimates on simulated and real datasets. It is the first method to reliably call gene-fusion events from long RNA reads. Sequencing the K562 transcriptome, we used AERON and found known as well as novel gene-fusion events.

## Introduction

Whole-transcriptome sequencing has become an important method in many research projects. Due to the revolution in short-read sequencing technologies and algorithm development, the detection of expressed RNAs in biological samples with thousands of cells [1] and even single cells [2] is nowadays done routinely. There are many important applications of such technologies, for example the detection of disease specific gene expression patterns [3] or the detection of gene fusion events in cancer cells [4], which opens the door for new therapeutic options and novel biological discoveries. However, short-read whole transcriptome sequencing has its weaknesses. RNA molecules can be thousands of nucleotides long and recent studies have revealed that more than 95% of multi-exonic genes undergo alternative splicing [5, 6]. Short-read sequencing thus has important limitations when it comes to the accurate quantification of gene isoform expression levels and the detection of gene fusion events [7]. Recent long read sequencing technologies, like developed by Pacific Biosciences (Pacbio) and Oxford Nanopore Technologies (ONT), have made significant progress in sequencing output per run at dramatically reduced costs. Thus, an important current challenge is to develop methods that can use long read RNA sequencing data for tasks where long reads overpower short reads, such as transcript quantification and gene fusion detection. Given their ability to cover large fractions of each transcript – and frequently complete transcripts – longer reads hold the promise of more accurate expression estimates. Although there is much potential in using long RNA reads for transcript quantification, limited work has been done in this area and, to our knowledge, there are presently no tools for detecting gene-fusion events from long RNA reads.

Modern bioinformatics tools designed for short read RNA sequencing such as Cufflinks [8], Kallisto [9] and Salmon [10] map short reads to a reference transcriptome and estimate abundances. Similarly, for fusion detection, algorithms such as TopHat-Fusion [11], SOAPfuse [12], MapSplice [13] and others align reads to a reference using a “splice-aware” aligner. They detect fusion events by considering reads overlapping two different genes [14]. All the above methods deal with the specifics of short-read RNA-seq protocols, but have not been adapted to cope with long read sequencing yet. They are able to correct for biases inherent to short-read protocols and address the fact that short reads often ambiguously align to several transcript isoforms of the same gene.

For long read analysis one approach is to cluster reads *de novo*, that is, without using a reference, based on sequence similarity between read sequences directly, *e.g.*, ToFU [15] or isONclust [16]. Each cluster should ideally contain reads originating from the same gene or transcript. Quantification can then be achieved by simply counting the number of reads in a cluster. But many genes have regions consisting of similar sequences. Hence, the reads generated from these similar regions might get clustered together which would result in erroneous quantification. Also pairwise read similarity computation can be time consuming.

An alternative approach is to use available reference sequences for the alignment of the long reads. However, long read alignment comes with its own challenges, and methods optimized for short-read data typically are not appropriate. Therefore specialized alignment packages have been developed to cope with the higher amount of sequencing errors of long-read technologies, *e.g.*, BLAT [17], BLASR [18] and Minimap2 [19]. Recent long read RNA analysis methods such as TALON [20] and Mandalorian [21] rely on these alignment programs to align long mRNA sequences against a reference genome. The quantification of a transcript is achieved by counting the number of reads overlapping a given transcript. Similarly, a recent study by Soneson *et al.* [22] used a pipeline that performed quantification by running Salmon on the alignments produced by Minimap2.

In all the above methods, the aligner finds a seed in the reference sequence and extends the alignment from the seeds. In case of multi-mapping reads, the aligner selects the primary alignment based on the alignment score. This makes the assignment of reads to a transcript biased towards how the primary alignment is selected [22]. Hence, if a read maps to multiple transcripts, which are highly similar to each other, the assignment becomes ambiguous.

Already in 2002, the concept of splicing graphs to represent possible transcripts of a gene has been introduced [23]. More recently, general purpose tools for working with sequence graphs have emerged [24–26]. Graphs with nodes representing nucleotide characters and edges representing adjacencies have successfully been used for variant calling [24–26], genome assembly [27–30], and short tandem repeat resolution [31]. Furthermore, alternative splicing events can be detected by aligning short reads to splicing graphs [32]. So far, to our knowledge, no algorithm has been proposed to use sequence graphs for long read transcript quantification and gene-fusion detection.

### Contribution

Here, we introduce AERON, an alignment based pipeline for quantification and detection of gene-fusion events using only long RNA reads. We adapt the sequence-to-graph alignment tool GraphAligner [33] to align reads generated from noisy long read technologies, such as ONT, to a reference transcriptome represented as gene-exon graphs. These graphs naturally allow to quantify gene and transcript expression based on the overlap with paths of known transcripts. We introduce the first long-read-specific gene-fusion detection algorithm. We tested AERON on three ONT datasets of varying coverage, including K562 data we generated for this study, and were able to achieve accurate quantification exceeding performance of current techniques. Prediction of fusions on simulated and real datasets illustrates the ability of AERON to discover known and novel fusion events directly from long read data. AERON is an open source software, which can be accessed via https://github.com/SchulzLab/Aeron (MIT licensed)

## Results

### A new approach for long read alignment and quantification

Here we present the AERON pipeline, which aligns long RNA reads to sequence graphs and reports the abundances of transcripts present in the sample. Figure 1 summarizes the approach. In a pre-processing step, a *gene-exon* graph is constructed for each gene of the genome (Fig. 1a). Each node in the graph for gene *g* represents an exon. If an exon is subject to alternative splicing in some of the transcripts, *e.g.* alternative donor or acceptor sites, the corresponding node is divided into sub-nodes. The 3’ end of a node is connected to the 5’ end of all the nodes downstream of it. The set of all gene-exon graphs represents the transcriptome. Each annotated transcript is uniquely represented as a path in one gene-exon graph. These graphs are constructed and augmented with the corresponding transcript paths as a preprocessing step, which we refer to as *indexing* (Fig. 1a,b).

**Figure 1.**
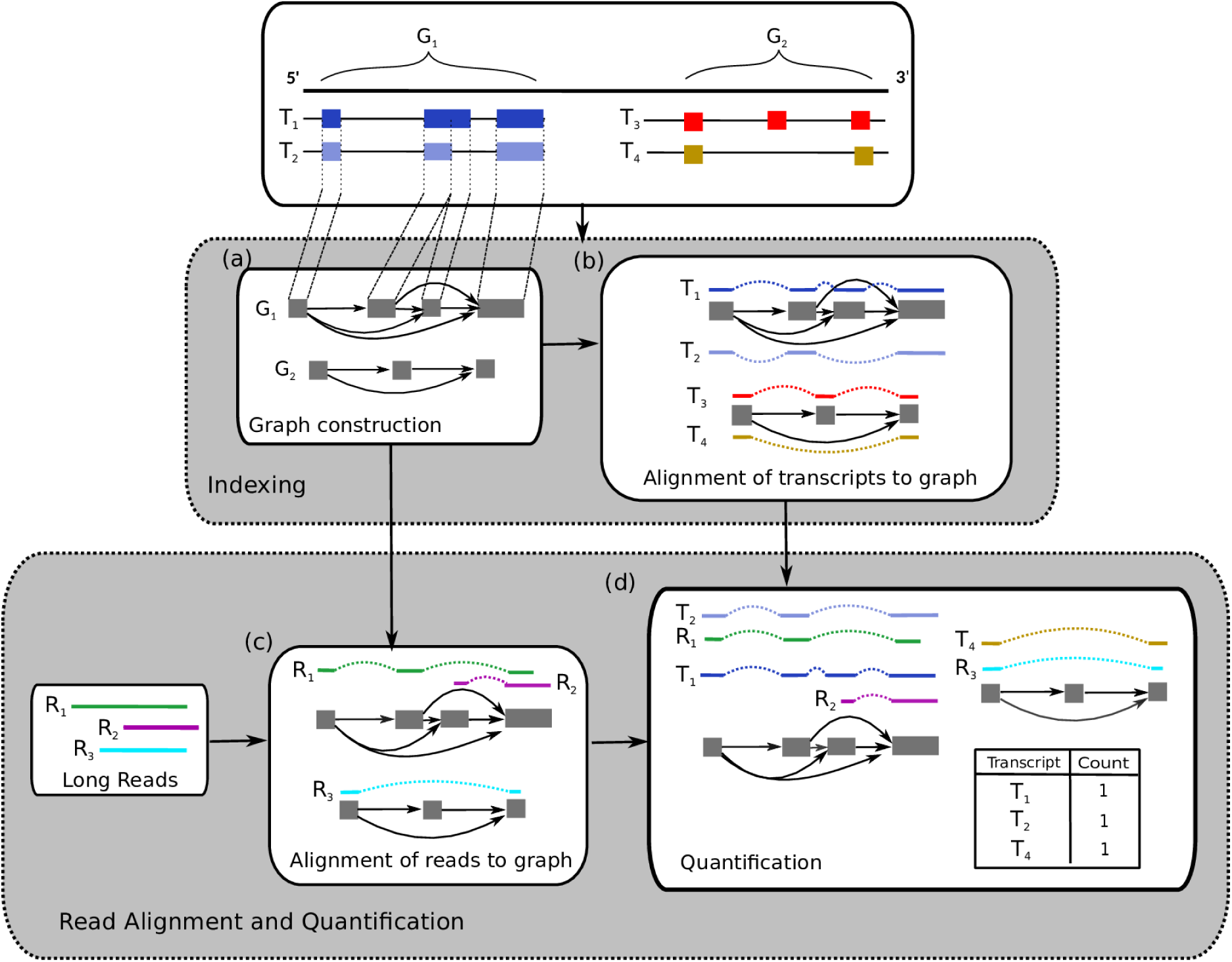
Workflow of the quantification step of AERON. Top white box depicts two genomic regions *G*_1_ and *G*_2_ consisting of four transcripts *T*_1_ and *T*_2_ in *G*_1_ and *T*_3_ and *T*_4_ in *G*_2_. The boxes within each transcripts depicts the exons present in the gene. The quantification consists of two steps -1) *Indexing:* this step begins with (a) graph construction where each exon of a gene serves as a node (grey box) and is split if the corresponding exon has an alternate donor/acceptor site. A node is connected, by an edge (solid arrows), to all nodes which are downstream of it. Graph construction is followed by (b) alignment of transcripts to the graph. 2) Index construction is followed by (c) alignment of reads to all the graphs. Alignments of step (b) and (c) are depicted by solid colored line and the dashed colored lines depicts the path followed by the transcript or the read. In the (d) quantification step, read and transcript alignment are compared. A read is assigned to a transcript if the path followed by the read is contained in the path followed by the transcript.

In the alignment step, a long read sequencing dataset is aligned to all gene-exon graphs (Fig. 1c) using the GraphAligner [33]. Finally, through comparison of read alignments with graph paths of annotated transcripts, expression values are computed (Fig. 1d). A transcript *t* is considered as expressed if for any read *r* the following conditions are met: 1) the E-value [34] of the alignment between *r* and the gene-exon graph is less than 1.0 and 2) at least 20% of the path covered by *r* is contained in the path traversed by *t*. The read *r* is assigned to transcript *t* with a score *s*, where *s* is the percentage of the path of *r* contained in *t*.

However, a read can – in principle – get aligned to two different transcripts with the same score. Earlier studies have shown that since sequencing is performed from the 3’ to the 5’ end, there is a certain bias towards the 3’ end of the transcripts [35], i.e, more reads are expected to come from a region near the 3’ end of a transcript. We observed the same effect as the density of reads was higher towards the 3’ end of the transcripts as compared to the 5’ end (Supp Fig.S1 and S2). Hence, in the scenario where a read is aligned to two transcripts with the same score, we assigned the read to the transcript whose 3’ end is closest to the 3’ end of the read. AERON assumes that each long read represents an individual transcript molecule.

The final quantification of transcript *t* is done by simply counting the number of reads assigned to it and converting the count into *Transcripts Per Million* (TPM) values (Fig. 1d). Gene-level quantification is achieved by summing up the TPM values for all the transcripts belonging to a gene.

### Comparison of sequencing protocols

Sequencing of a transcriptome generally involves synthesizing the complementary DNA (cDNA) and amplifying the cDNA using PCR. An alternative protocol is to directly synthesize the single stranded RNA (DirectRNA) [36]. To measure the performance of AERON on data generated from the above two protocols, we downloaded two ONT datasets from the NA12878 cell line [37]. The first contained 15M sequences obtained using the cDNA protocol and the second 10M sequences obtained using the DirectRNA protocol. We ran AERON on these two datasets separately. As expected, a higher density of reads in both datasets aligned closer to the 3’ end of the transcript (Supp. Fig. S2). This 3’ bias was stronger in alignments of *DirectRNA* reads as compared to reads from the *cDNA* dataset.

Further, we computed the expression levels of all known Ensembl genes and transcripts (version 92) for both datasets. Out of 58,336 annotated Ensembl genes, 28,584 and 28,021 genes were identified using the *cDNA* and *DirecRNA* dataset, respectively. 25,823 genes were commonly identified in both. Similarly, out of 203,675 annotated transcripts, 102,748 and 107,030 transcripts were identified from the *DirectRNA* and *cDNA* subset, respectively. 85,697 transcripts were commonly detected in both. Figure 2 shows a comparison between the protocols at gene and transcript level. We found gene expression levels to be highly correlated across the protocols (Spearman r=0.90). Expectedly, at the transcript level, the correlation between the expression estimates from the two protocol was lower, but still in reasonable agreement (Spearman r=0.68, Figure 2b). Further, we randomly divided the reads of NA12878 dataset into three subsets and ran AERON on the three subsets seperately with default parameters. We calculated the gene level and transcript level expression for the three subsets. We found both the gene level and transcript level estimates to be highly correlated to each other (Supp. Fig. S2 and S3). Based on the above experiments we were able to assert the fact that ONT data quantified using AERON produces reproducible quantification.

**Figure 2.**
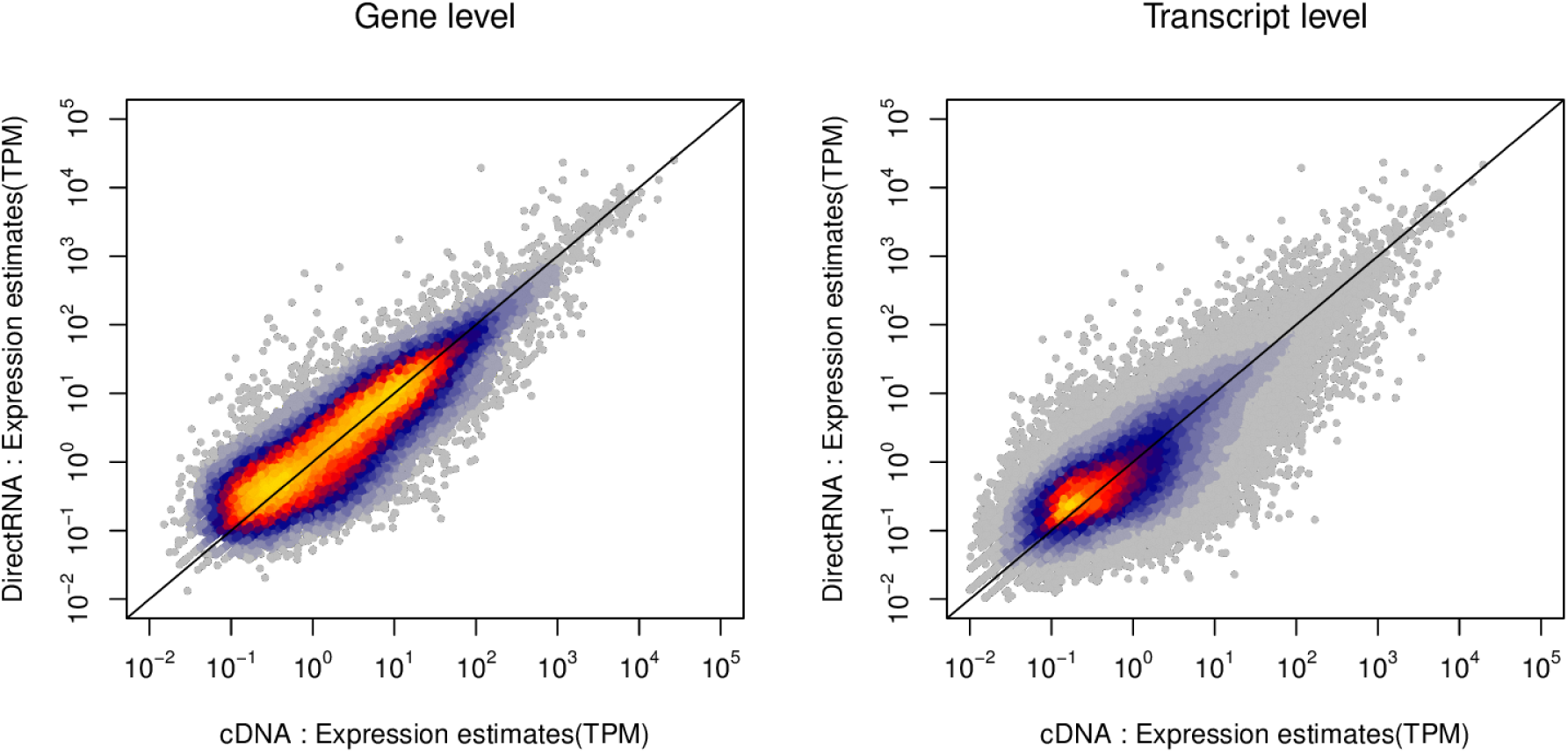
Performance of AERON on datasets obtained using two different ONT protocols. The scatterplot compares the expression estimated (TPM values) obtained from the RNA-seq sequences from cRNA (x-axis) against the expression estimated obtained from RNA-seq sequences from native RNA (y-axis) at gene and transcript level.

### Comparison with other methods

To further test the performance of AERON, we generated 2M ONT reads from the K562 cell line. We aligned and quantified the NA12878 and K562 reads against the human Ensembl transcriptome. In the K562 dataset, more than 80% of the reads were aligned with a median length of 398. For NA12878, 96% of the reads were aligned to the transcripts with a median length of 776. As expected, many of the unaligned reads were either short in length or contained many low quality bases, which suggests they contain too many errors to determine their origin correctly (Supp. Fig. S5).

In order to compare estimates from AERON and Minimap2 [19] using our long read datasets, we computed an additional estimate of gene and transcript expression for both cell lines from short read Illumina data using Salmon [10], see Methods. Taking the Salmon estimates as reference allowed us to calculate Spearman correlation and absolute error values (MARD scores, see Methods). In these comparisons we considered all genes and transcripts present in the human genome according to ENSEMBL annotation (v92). We summarized the results in Table 1. For the K562 dataset, AERON is able to assign approximately 50% more reads to genes as compared to Minimap2. For the complete NA12878 dataset, AERON is able to assign 3% more reads than Minimap2. AERON shows better correlation and error rates, i.e, lower MARD scores than Minimap2 when compared to short read gene expression estimates for both datasets.

**Table 1.** Spearman correlation and MARD between Transcripts Per Million (TPM) at gene level obtained from AERON/Minimap2 using Oxford Nanopore Sequencing (ONT) data and TPM at gene level obtained from Salmon using Illumina data. The size of the dataset in millions(M) is depicted in brackets next to the name.

| Dataset | AERON |  |  | Minimap2 |  |  |
| --- | --- | --- | --- | --- | --- | --- |
|  | #Mapped | Correlation | MARD | #Mapped | Correlation | MARD |
| K562 (2.7M) | 2,1M | **0.833** | **0.350** | 1M | 0.704 | 0.428 |
| NA12878 (25M) | 23,9M | **0.822** | **0.349** | 23,2M | 0.778 | 0.37 |

Figure 3 shows a scatterplot of gene expression estimates of AERON (Fig. 3a,c) and Minimap2 (Fig. 3b,d) against the short read estimates (y-axis). For this plot, we only take the genes which had non-zero expression estimate either by AERON or by Minimap2. We observe that, for both datasets, AERON is able to assign reads to the correct gene, resulting in a majority of points being located close to the diagonal. Moreover, we see that AERON is able to detect more genes in lower range of expression (1-5 TPM) as compared to Minimap2. Similar behavior was observed at transcript level, but the correlation values between the expression estimates from long read data and estimates from short reads were lower (Supp. table S1). Beyond challenges to do transcript quantification from short reads, a possible reason for this might be the presence of a high number of short transcripts against which no reads are mapped. To test this hypothesis, we filtered out all the transcripts below a length cutoff from our correlation analysis experiment. As expected, when we remove short transcripts, the correlation improves for both datasets supporting our hypothesis. For a range of transcript length cutoff of 0-10000bps, the correlation improved from 0.29 to 0.64 for K562 cell line and 0.27 to 0.74 for NA12878 dataset(Supp. Fig. S6).

**Figure 3.**
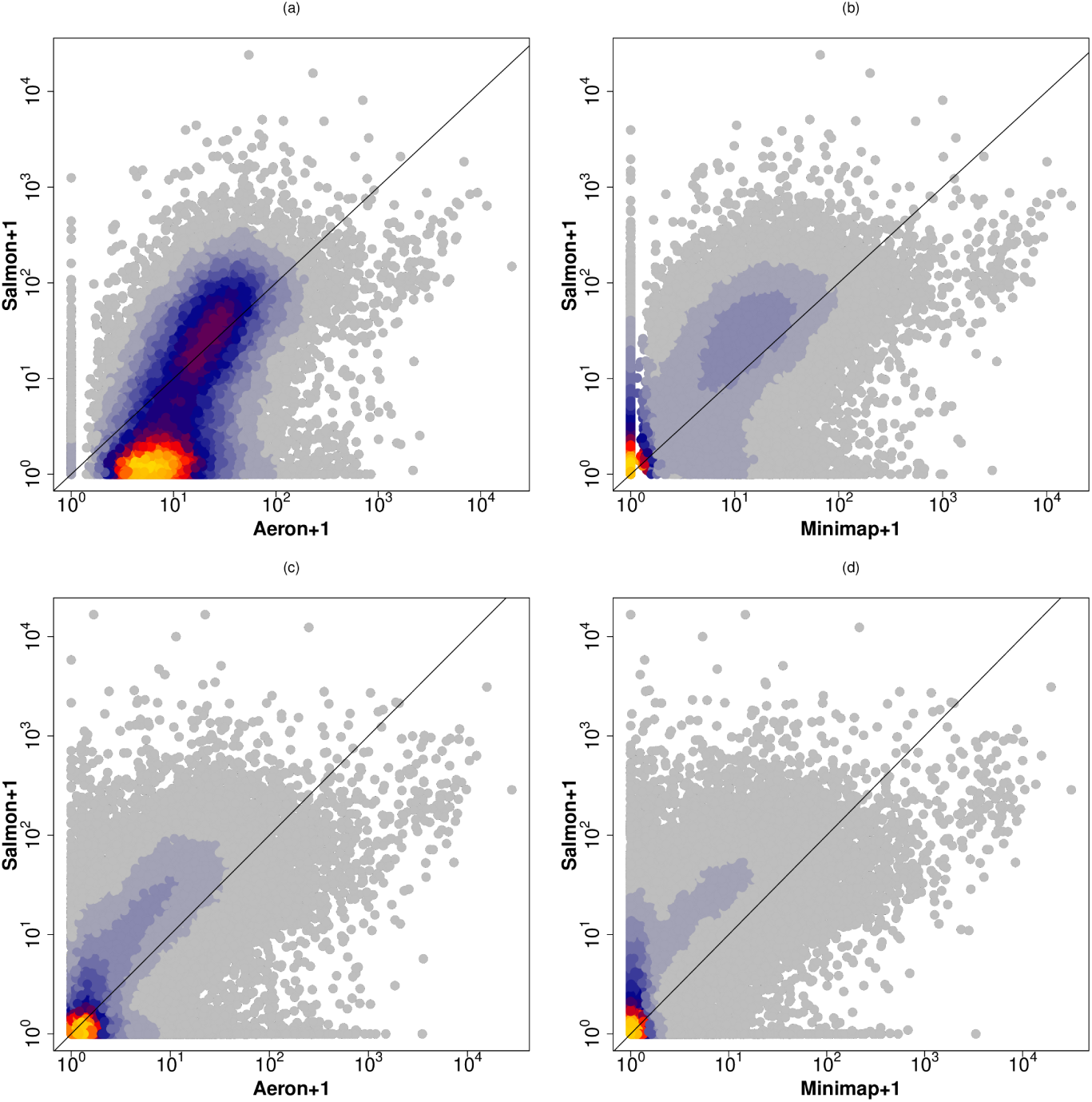
Comparison of gene level expression estimates obtained from AERON (a and c) against the expression estimates obtained from Minimap2 (b and d) for the two datasets - K562 (a and b) and NA12878 (c and d). The expression estimated using long reads are the shown in the x-axis and the estimates obtained using short reads are shown in the y-axis. For clarity, expression estimates have been log transformed with a base 10.

### Fusion simulation

The previous experiments underline that AERON is able to align noisy long RNA reads to transcripts accurately. Motivated by a lack of approaches that enable fusion-gene detection with long reads and the high relevance of fusion genes, we implemented a fusion detection method. Figure 4 shows an overview of the pipeline. There are three main steps in the fusion detection pipeline: First, partial alignments from the reads provide *tentative fusions*. Next, the reads are re-aligned to *fusion graphs* derived from the tentative fusions. Finally, the alignments to the fusion graphs are compared with alignments to gene-exon graphs to provide a *fusion score* for each read. Each of the steps produces a list of fusion event candidates, with the later steps filtering out candidates from the previous steps. The methods section describes the individual steps in more detail.

**Figure 4.**
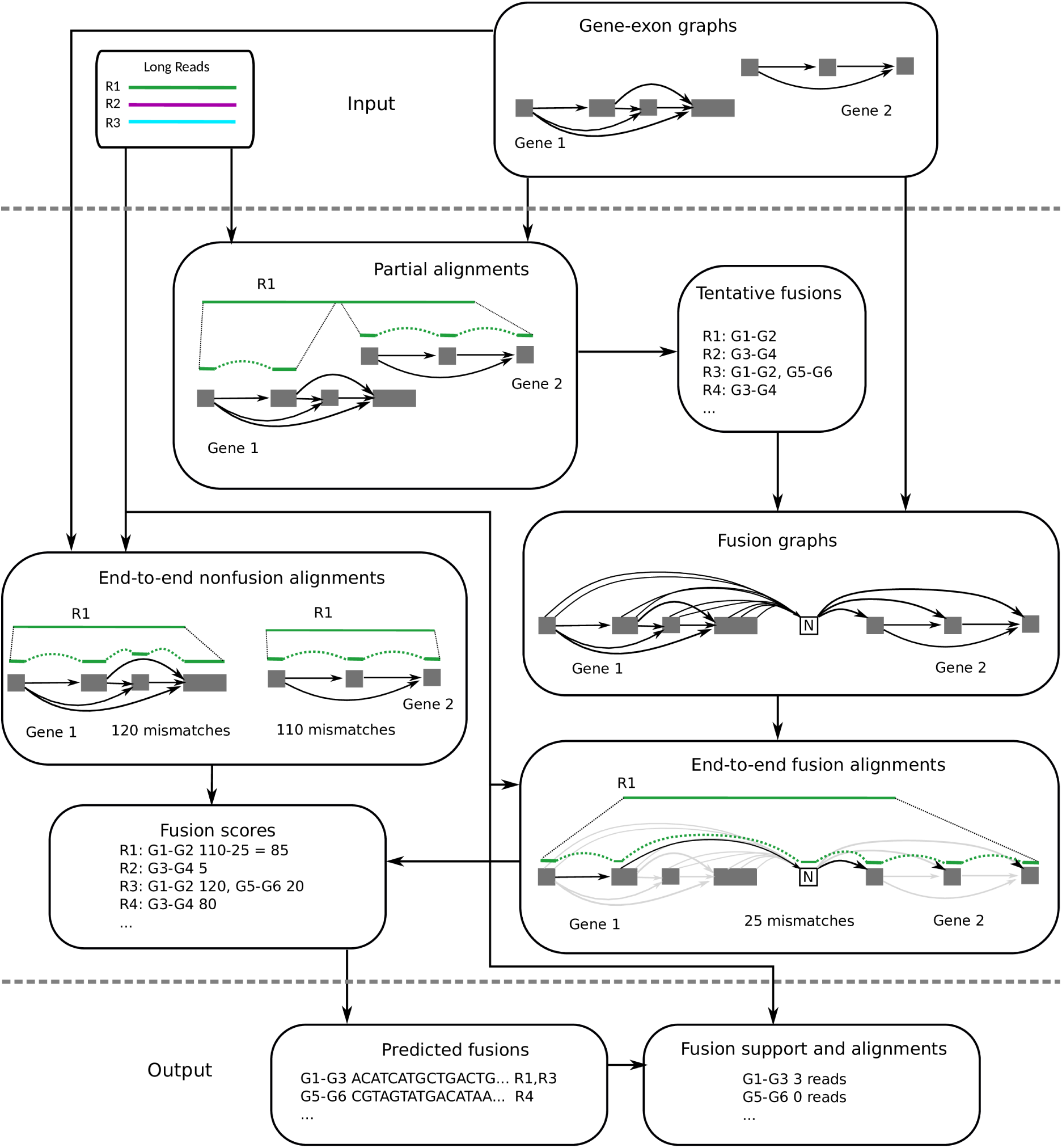
Workflow of the fusion detection step of AERON. Partial alignment: reads are aligned to the gene-exon graphs. All secondary alignments are kept and the read may have alignments to different genes. Tentative fusions: whenever a read has a pair of alignments that end within 20 bp of each other in the read, the read votes for a fusion between the two genes. A read may vote for multiple tentative fusions. Fusion graphs: each tentative fusion induces a fusion graph, where the two genes are connected with a crossover node (N). End-to-end fusion alignments: the reads are aligned to the fusion graphs. Global alignment is used to align the read from start to end. End-to-end nonfusion alignments: the reads are aligned to the individual gene-exon graphs globally. Fusion score: the score difference between the fusion alignment and the nonfusion alignment defines a fusion score. Predicted fusions: the alignments are filtered based on the fusion score. The graph sequence along the alignment is taken as the predicted fusion transcript. Fusion support and alignments: all reads are aligned to the reference transcripts and the predicted fusion transcripts with Minimap2. A read supports a fusion if its primary alignment covers the fusion breakpoint with at least 150 base pairs on both sides.

We first assessed the performance of our fusion detection approach in a simulation study. We generated fusion events of different “lengths”, where the length refers to the amount of sequence from both genes. For example a fusion event with length 200 bp contains 200 base pairs from both genes and has a total length of 400 bp. Events were simulated in 9 length groups, from 100-200 bp, 200-300 bp, and so on until 900-1000 bp. For each of these length ranges, 50 fusions were generated, where a pair of transcripts was selected randomly for each fusion. Then, a random substring of each transcript corresponding to the length of the fusion was selected, and the substrings were concatenated to build the fusion transcript. The reads were simulated at 10× coverage from all simulated fusion and reference transcripts. The fusion detection pipeline was then ran on the simulated reads.

The left part of Fig. 5 shows the precision-recall curve at varying fusion score cutoffs for different fusion sizes. We see that 100-400 base pair fusions (bottom) are hard to detect with any fusion score cutoff, and the recall saturates at 15%. The high error rate and short length of the reads stops them from being aligned in the tentative fusion phase, which prevents the pipeline from detecting the fusions. However, 400-700 base pair fusions (middle) are detected, and the recall reaches up to 87%. For the longer fusions of 700-1000 base pairs (top), recall reaches up to 95% and precision around 80%. Based on the curves, we chose 200 as the default fusion score cutoff, as that achieves a precision of 78% and recall of 90% for large fusions.

**Figure 5.**
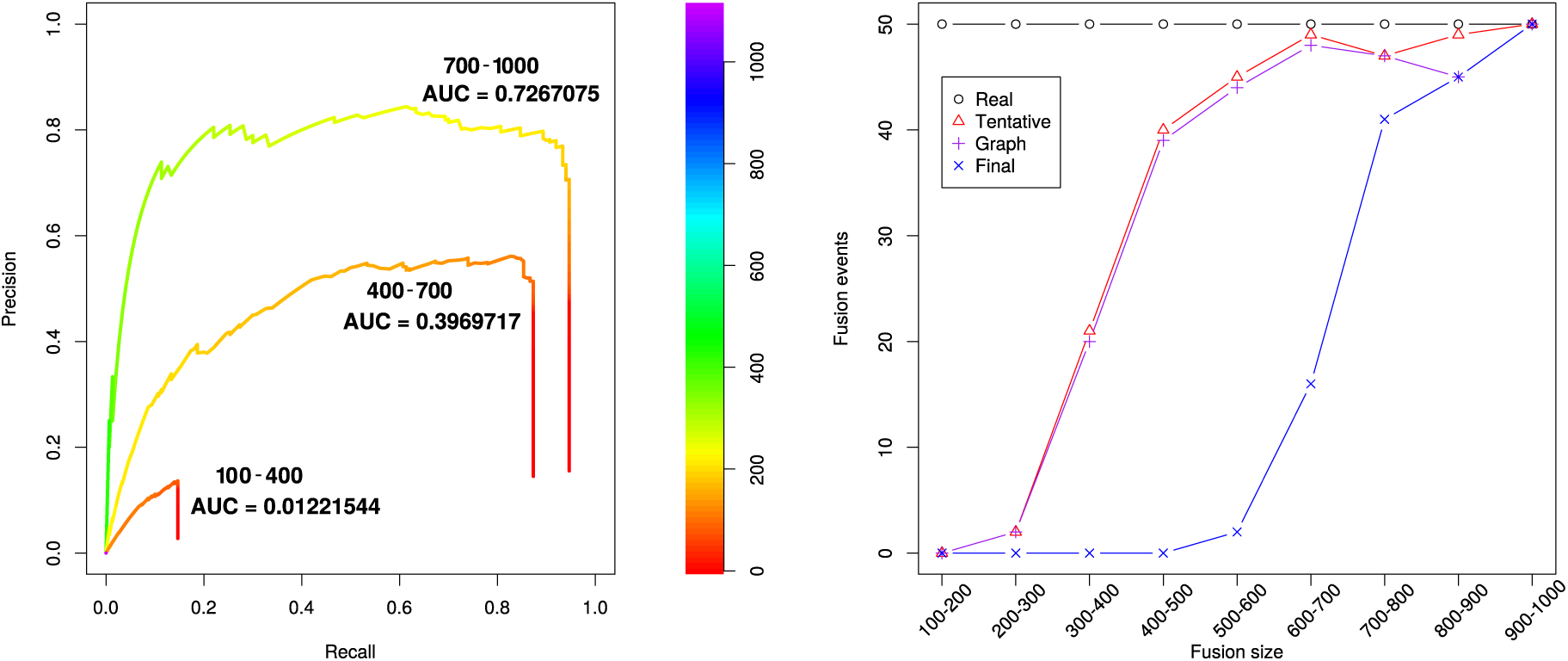
Left: Precision-recall curves for fusion event detection with simulated data for fusions of 100-400 base pairs (bottom), 400-700 base pairs (middle) and 700-1000 base pairs (top). Both precision and recall improve for longer fusions. The parameter varied is the fusion score cutoff. Right: number of detected true fusion events per fusion size with simulated data. The curves show the number of simulated fusions (Real) and fusions detected at different parts of the pipeline: *tentative fusions* (Tentative), after fusion graph alignment (Graph), and after filtering for fusion score (Final). The number of total false positives is 28696 for Tentative, 49 for Graph and 20 for Final.

The right part of Figure 5 shows the number of detected fusions (true positives) as a function of fusion length at different phases of the fusion detection pipeline. These experiments also show that our three-step approach progressively removes false positive fusion events: while there are 28,696 false positive predictions in the tentative step, this number reduces to 49 in the graph step and further down to 20 in the final step. We see that shorter fusions are not detected even in the tentative fusion phase; this is most likely due to the high error rate preventing short alignments from being found. Once the fusion size grows above 400 bp, the set of tentative fusions contain most of the fusion events. Importantly, the fusion graph approach removes only a small fraction of true fusions, while removing almost all false positives from the tentative fusions. The fusion score cutoff further removes more false positives, but at the cost of removing shorter true positives as well.

### Fusion detection on real data

We ran the fusion detection pipeline on the two datasets: NA12878 and K562. The NA12878 functions as a control, as we do not expect to see any fusions in that dataset. K562, on the other hand, is a highly rearranged cancer cell line [38] with a known BCR-ABL1 fusion.

We used predictions which were supported by at least two reads, resulting in 25 events for K562 and 24 for NA12878. We then further manually curated the predictions in three steps. First, we removed all fusions involving a mitochondrial gene and removing fusions between a gene and its own pseudogene, leaving 16 events for K562 and 14 for NA12878. Then we used IGV [39] to visually inspect the alignments of the reads to the predicted fusion transcripts generated by the pipeline, and discarded events where the alignments to one of the genes seemed noticeably worse than the other or the read coverage over the breakpoint was noticeably less than for either of the genes. Supplementary Figure S7 shows an example of an event that was rejected based on the IGV visualization. After IGV visualization, 15 events were left for K562 and 9 for NA12878. Since some transcripts can have similar sequences, the high error rate of the reads can cause a false positive fusion prediction between the two transcripts due to sequence similarity. In this case, the predicted fusion transcript should be similar to a reference transcript, because it reflects a reference-guided consensus between the long reads. Hence, the average error rate of the fusion transcript is much lower than the input reads and therefore easier to align with traditional methods. In order to detect transcripts not annotated in the Ensembl release, we used the BLAST webserver to align the predicted fusion transcripts to the human reference genome (GRCh38.p12) and transcriptome (NCBI Homo sapiens Annotation Release 109). We discarded events that mapped to an existing transcript including the fusion breakpoint and retained 8 events for K562 and 2 events for NA12878.

The two predicted fusion events for NA12878 are shown in Table 2. Supplementary figures S8 and S9 show corresponding IGV screenshots of the transcripts and their reads.

**Table 2.**
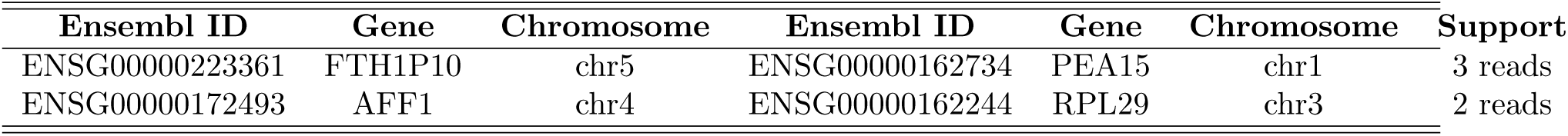
Predicted fusion events for NA12878. The predicted fusion transcripts do not have BLAST hits that cover the fusion breakpoint. The first 6 columns describe the two genes involved in the fusion. The column “Support” counts the number of reads whose primary alignment covers the fusion breakpoint and 150bp from both sides of it.

| Ensembl ID | Gene | Chromosome | Ensembl ID | Gene | Chromosome | Support |
| --- | --- | --- | --- | --- | --- | --- |
| ENSG00000223361 | FTH1P10 | chr5 | ENSG00000162734 | PEA15 | chr1 | 3 reads |
| ENSG00000172493 | AFF1 | chr4 | ENSG00000162244 | RPL29 | chr3 | 2 reads |

Table 3 shows the eight predicted fusion events for K562 including the well-known BCR-ABL1 fusion event (Fig. 6). Four of the eight predicted fusion events have been reported in literature before. For the BCR-ABL1 fusion, the TEN1-CDK3 read-through and BMS1P4-AGAP5 read-through events, the AERON predictions mapped to the existing annotations. The NOS3-PRKN predicted fusion mapped to a transcript variant of the NOS3 gene, while the ARPC4-TTLL3 predicted fusion mapped to a transcript variant of ARPC4. The PRIM1-NACA predicted fusion occurred with fusions across two different breakpoints, which the pipeline considers separate events. The HBG2-HBG1 predicted fusion has a very high read support, and the alignment of the predicted transcript to the reference is consistent with an inverted duplication (see Supplementary Figure S8) occurring in the region.

**Table 3.** Predicted fusion events for K562. The TEN1-CDK3 and BMS1P1-AGAP4 were reported earlier as read-through events. The first 6 columns describe the two genes involved in the fusion. The column “Support” counts the number of reads whose primary alignment covers the fusion breakpoint and 150bp from both sides of it.

| Ensembl ID | Gene | Chromosome | Ensembl ID | Gene | Chromosome | Support | Known |
| --- | --- | --- | --- | --- | --- | --- | --- |
| ENSG00000196565 | HBG2 | chr11 | ENSG00000213934 | HBG1 | chr11 | 89 reads | [40] |
| ENSG00000186716 | BCR | chr22 | ENSG00000097007 | ABL | chr9 | 2 reads | [41] |
| ENSG00000257949 | TEN1 | chr17 | ENSG00000250506 | CDK3 | chr17 | 2 reads | [42] |
| ENSG00000204177 | BMS1P1 | chr10 | ENSG00000188234 | AGAP4 | chr10 | 2 reads | [43] |
| ENSG00000241553 | ARPC4 | chr3 | ENSG00000214021 | TTLL3 | chr3 | 4 reads | – |
| ENSG00000198056 | PRIM1 | chr12 | ENSG00000196531 | NACA | chr12 | 3 reads | – |
| ENSG00000198056 | PRIM1 | chr12 | ENSG00000196531 | NACA | chr12 | 3 reads | – |
| ENSG00000164867 | NOS3 | chr7 | ENSG00000185345 | PRKN | chr6 | 2 reads | – |

**Figure 6.**
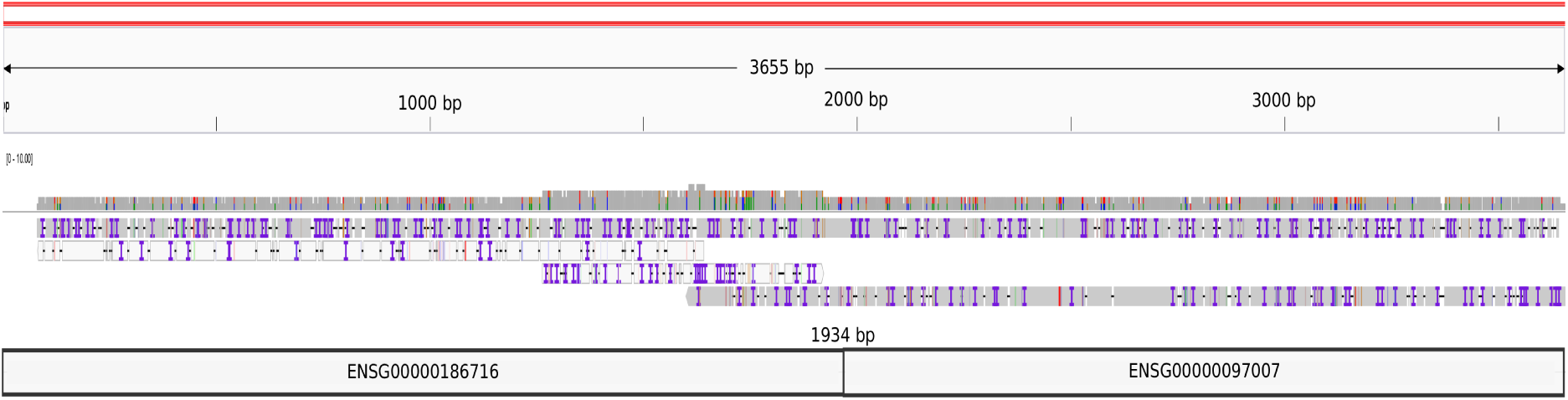
The coverage plot and the alignment of reads against the 3655bps long BCR-ABL1 fusion transcript. The fusion breakpoint was found to be at the position 1934. The image was generated using Integrated Genomics Viewer(IGV).

## Discussion

In this work, we describe AERON, a novel approach to quantify transcripts from long reads. AERON comes with the first long-read specific fusion detection method. We use GraphAligner [33], a fast sequence-to-graph alignment method, to align ONT reads to a reference transcriptome and find better alignments as compared to Minimap2, which is used as part of previous state-of-the-art quantification pipelines. We assign reads to transcripts based on the position of the read mapping within the transcriptome and the fraction of read sequence contained in the transcript sequence. We tested AERON on two different datasets of varying coverage and compared results to expression estimates derived from Minimap2 alignments of the reads to the transcripts. We compared the expression estimates from these two methods (AERON and Minimap2) against the estimates generated from short read data. We found that expression estimates from AERON had a better correlation as compared to estimates from Minimap2. We also show that AERON does not depend on the sequencing protocol used to generate the reads. This was evident from the high correlation between the AERON runs on reads generated from the cDNA and the reads generated from the native RNA.

Although we show improvement over the existing methods, there are remaining limitations. We find the gene-level quantification to be much better compared to the transcript level quantification. We have shown that the presence of short-length transcripts has an effect on the assignment of reads to the correct transcript, which is further complicated by highly similar transcripts. A significant number of transcripts are truncated versions of longer transcripts. In cases like these, it is difficult to decide whether a read has originated from the truncated transcript or is it a part of the longer transcript. This problem is further aggravated if the read originating from them does not overlap completely with either of the transcripts.

Soneson *et al.* [22] recently proposed an alternate approach where the EM based algorithm of Salmon was run on alignments produced with Minimap2 as input. We tested the suggested approach on both the datasets and found no major effect on the transcript level expression estimate (Supp.tab S3). The comparatively low correlation between estimates from short read data with estimates from the long read data at transcript level may be explained by the higher sequencing depth of short read datasets. This problem is mitigated to a certain extent in the gene-level quantification, where we add up the expression estimates of all transcripts belonging to a gene. Long read sequencing suffers from lack of sequencing depth, due to which many transcripts are either not identified or misidentified.

Furthermore, while our approach is able to align long error-prone reads, the analyzed data sets also contained relatively short reads, which are challenging to align given the high error rate. One way to potentially rescue such reads could be error correction. GraphAligner can perform error correction of long reads based on alignment of long reads against a De Bruijn graph generated from short reads. The process has been tested on genomic reads. Currently, no RNA-seq specific error correction tools are available, which could be integrated in our pipeline. A recent study by Lime *et al*. 2019 [44], shows that long read DNA correction methods improve analysis of long read RNA data, but have a negative effect on detecting diverse transcript isoforms and alternative splice sites. Therefore, we did not consider such approaches here, but this would be an interesting direction for further research.

We have also presented a novel fusion detection pipeline based on aligning reads to a fusion graph between two genes, filling an important gap in the landscape of tools for the analysis of long read RNA-seq data. We tested this on simulated data of different fusion lengths and real data from the K562 cell line. Experiments with simulated data show that the accuracy depends on the fusion length and fusions which contain at least 700 base pairs from both genes are detected reliably. Experiments with real data on the K562 cell line recovered four known events, including the BCR-ABL1 fusion, and suggested four novel events. In the future, we plan to automate further curation steps which we performed manually for this study. As the simulation experiments show that long fusion events are recovered with high recall, it is unlikely that the sample has other long fusions with significant expression.

## Methods

### K562 cell culture

K562 suspension cells were maintained in RPMI plus 10% FBS and contains 1% of Gibco Penicillin Streptomycin Solution 10,000 U/mL cat no: 15140122, culture in 37C, 5% CO2 incubator.

### RNA isolation

Total RNA was isolated from K562 cell line using TRIzol reagent (ThermoFisher Scientific) according to the manufacturer’s instructions. Total RNA concentration was measured at 1:10 dilution using a Qubit 3.0 Fluorometer and Qubit HS RNA Assay Kit (ThermoFisher Scientific). RNA quality and RNA integrity (RIN) were evaluated using a NanoDrop 2000 spectrophotometer (ThermoFisher Scientific) and Agilent 4200 TapeStation system with the RNA screentape assay kit (Agilent).

### Nanopore cDNA library generation and sequencing

Poly(A)+ RNA was enriched from 25*µ*g total RNA using the Dynabeads mRNA Purification Kit (ThermoFisher Scientific) according to manufacturer’s instructions. The K562 cDNA libraries were generated from 5ng of poly(A)+ RNA using the PCR-cDNA kit, SQK-PCS109 (Oxford Nanopore Technologies) according to manufacturer’s instructions. In brief, complementary strand synthesis and strand switching were performed on input full-length poly(A)+ RNA using kit-supplied oligonucleotides, followed by 14-cycle PCR amplification using primers containing 5’ tags to generate double-stranded cDNA library, followed by a ligase-free attachment of rapid sequencing adaptors. Final library was loaded onto a FLO-MIN106, R9.4 flowcell (Oxford Nanopore Technologies) and sequenced on a MinION Mk1B device with MinKNOW v18.12.4 for 48 hours. Post-run base calling was performed using Albacore 2.3.3.

### Index construction

The splicing graph generated from the reference sequence and the paths in the graph followed by all the annotated (according to ENSEMBL annotation-v92) transcripts in the splicing graph forms the index for transcript quantification and fusion detection. These steps are performed only once and can be re-used for multiple input datasets.

#### Graph construction

The set of all possible transcripts of a gene can be expressed as a splicing graph, first introduced by Heber *et al.*, 2002 [23]. In general, a splicing graph is a Directed Acyclic Graph (DAG) where nodes represent the splicing sites of a given gene and edges represents exons and introns between the sites. In this work, we slightly modify the splicing graph construction and name the graph as *gene-exon* graph, which we later align long reads to. Below, we first describe the construction of the graph from genome sequence.

We formally define a *DNA* sequence as a string consisting of characters from the alphabet S = {*A, C, G, T*}. A gene *g* is a *DNA* sequence consisting of exons and introns. We term a base within the gene *g* as *exonic base* if it is overlapped by at least one exon belonging to *g*. Otherwise, the base is termed as *intronic base*. A position in *g* is termed as a *border*, if it serves as a boundary of an exon. We classify borders into 5’ borders (*a*), which are positions of the 5’ end of an exon, and 3’ borders (*b*), which are the position of the 3’ end of an exon. Each exon *x* can be characterized a pair these two border positions, i.e, *x* = (*a, b*). We refer to a list of all 5’ borders and 3’ borders as *border list ρ*. We can divide the border list into two sub lists namely the *acceptor list α*, which contains all 5’ borders and the *donor list δ*, which contains all 3’ borders of *g*. Members of each list and sub lists are ordered increasingly. For instance, let exon *x*_1_ = (5, 10) and *x*_2_ = (15, 20) be two exons of the same gene *g* of length 20 bps, then *ρ*(*g*) = [5, 10, 15, 20], *α*(*g*) = [5, 15] and *δ*(*g*) = [10, 20].

The *gene-exon* graph of a gene *g* is defined as a directed acyclic graph *G*_*g*_ = (*V, E*)_*g*_. Each vertex of our *gene*-*exon* graph represents an exon present in the gene and the edges between vertices corresponds to all the possible splicing events occurring in the gene. If an exon has alternate splice sites, then the vertex corresponding to the exon is split into multiple sub-vertices. The split happens at the position of the alternate site. This concept is formalized in the following way. Each vertex *V* of the graph is considered as a substring between two consecutive borders of *g*. Let *i* ∈ *ρ*(*g*) and *j* ∈ *ρ*(*g*) be two consecutive borders in *g* where *i* < *j*. Then the vertex *v*_*ij*_ ∈ *V* is created using the following function.

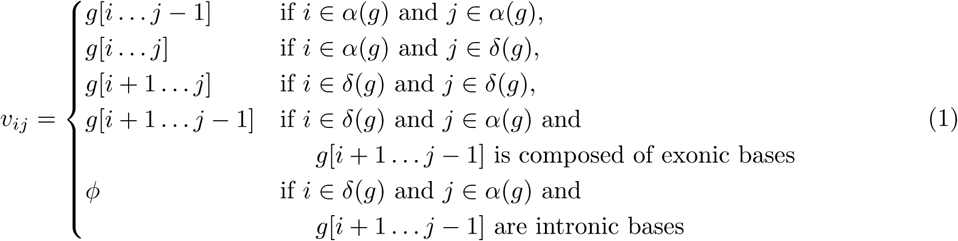

where *v*_*ij*_ = *ϕ* represents an null vertex. In a *gene*-*exon* graph, there should always be a path in the graph which represents all possible alternative splicing events. For instance, let there be three vertices *v*_12_,*v*_34_ and *v*_56_ corresponding to three exons *x*_12_, *x*_34_ and *x*_56_ respectively. If a transcript is formed by splicing out *x*_34_ and using only *x*_12_ and *x*_56_, then this event should be represented in the graph as a path passing through vertices *v*_12_ and *v*_56_. Hence, in the graph, given two non-null vertices *v*_*ii*′_ and *v*_*jj*′_, an edge *e*_*i*′*j*_ is created between all vertices with *i′* < *j*. Figure 7 provides a visual representation of the graph construction process and the implementation details are given in Supplementary Material Section 1.

**Figure 7.**
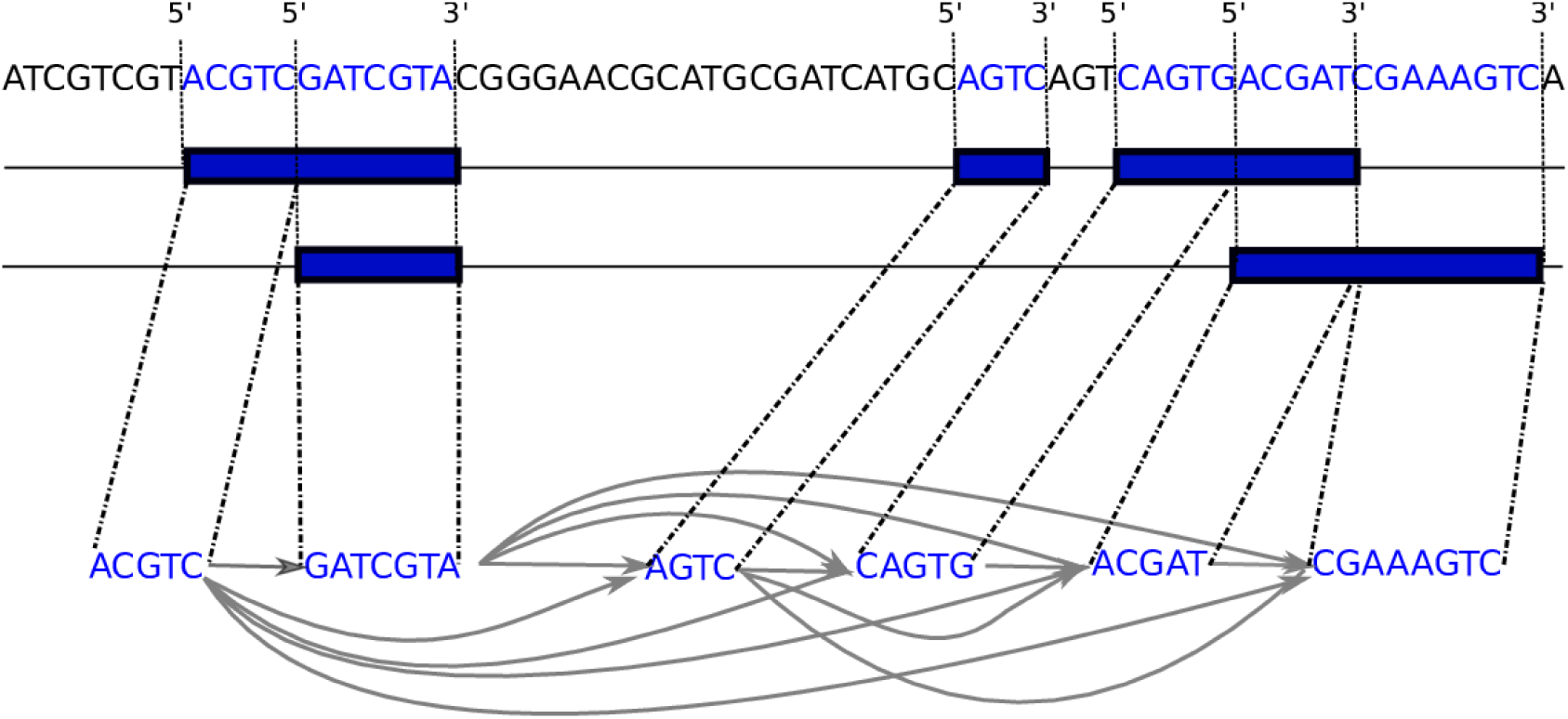
Construction of gene-exon graph from reference sequence. The exons are represented as blue boxes and its corresponding sequences are highlighted in blue. A border position is labelled as 5′ if it is the 5’ end of an exon or 3′ if the position is the 3’ end of an exon. Nodes of the graph are sub-sequences between two consecutive border positions. The 3’ end of the node is connected by an edge to the 5’ end of all the nodes downstream of it.

#### Sequence-to-graph alignment

We need to be able to align a sequence to a gene-exon graph for two purposes: first, for aligning known transcripts to the graph during index construction and, second, for aligning reads to the graph for quantification and fusion detection. The goal of this step is to find the *path* in the graph for each read sequence with the lowest edit distance.

Given a gene-exon graph *G*_*g*_ = (*V, E*), we define a path as a list of nodes *p* = *v*_1_, *v*_2_, …, *v*_*n*_ where *v*_*i*_ ∈ *V* and (*v*_*i*_, *v*_*i*+1_) ∈ *E*. The *path sequence* of a path is the concatenation of the node labels of *p*. Given a read sequence and a graph, the *sequence-to-graph alignment* problem is to find the path in the graph with the smallest edit distance between the path sequence and the read sequence. We use GraphAligner for aligning the reads [33]. Briefly, GraphAligner is a seed-and-extend aligner that finds maximal exact matches (MEMs) [45] between the read and the node sequences and then extends them with a bit-parallel dynamic programming algorithm [46]. This produces a set of *alignments*, where each alignment contains a path, edit distance, an E-value [34] and the start and end positions in the read. The parameters used by GraphAligner are the maximum number of MEMs to extend and the minimum MEM length.

#### Alignment of transcripts to graphs

Given a genome consisting of *n* genes and *m* transcripts, we begin by constructing a *gene*-*exon* graph *G*_*i*_ for each gene in the genome and add it to a graph set *U* = {*G*_1_, *G*_2_, ….*G*_*n*_}. We align the *m* reference transcripts to all the *n* graphs present in set *U*. For each transcript *t*, we obtain the path which has the minimal edit distance to the transcript. Since the nodes of a graph are composed of exonic sequences from the genomic regions, the best possible path traversed by transcript *t* is the gene-exon graph of the gene from which *t* originates. Hence, each transcript will only have one path associated with it. We collect all such paths in set *P* = {*p*_1_, *p*_2_, …, *p*_*m*_} and name the set as *transcript-path* set.

### Assignment of reads to transcripts and quantification

Consider the task of aligning a long read set *R* = {*r*_1_, *r*_2_, …, *r*_*k*_} consisting of *k* reads. In principle we align all reads in *R* to all the graphs in the set *U*. We only consider alignments with an *E-value* [34] *below 1. In case a read r* ∈ *R* has multiple alignments meeting this criterion, we select the longest alignment. The first condition removes poor alignments between the read and the graph and the second condition ensures that the read is not mapped to multiple locations. The E-value [34] of the alignment between a query and a database describes the expected number of spurious alignments that are at least as good as the alignment. The E-value is calculated with the formula:

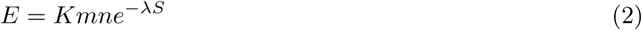

where *E* is the expected number of spurious hits, *K* and *λ* are parameters that depend on the scoring scheme, *S* is the alignment score, *m* is the database size in base pairs and *n* is the query size in base pairs. The original formula by Karlin and Altschul is defined with a database of linear sequences instead of graphs. We use the number of base pairs in the graph as the database size. The *K* and *λ* were chosen to correspond to a scoring scheme with match score +1 and mismatch cost -2.

Once the reads and the transcripts have been aligned, we extract only the path of the alignment and discard the base pair level alignments. That is, each read will be treated as a string *q*_*r*_ = *v*_1_*v*_2_…*v*_*n*_ where *v*_*i*_ ∈ *V* and the differences between the read sequence and the path sequence are ignored. We then compare *q*_*r*_ to all the paths in the *transcript path* set. We say that a read *q*_*r*_ belongs to a transcript *p*_*i*_ if there is a node that both paths include, that is, ∃_*v*_ : *v* ∈ *q*_*r*_ ∧ *v* ∈ *p*_*i*_. For each read path *q*_*r*_ belonging to transcript path *t*_*i*_, we align the *q*_*r*_ and *t*_*i*_ to each others using the Needleman-Wunsch algorithm [47] with nodes weighted according to the amount of sequence in them. We then define the *overlap score* as the fraction of matches in the alignment over the read length. The overlap score represents the fraction of the read which matches the given transcript. If the *overlap score* is above 20%, then *q*_*r*_ is said to be aligned to *t*_*i*_.

A read may get aligned to multiple transcripts. We assign each read to the transcript with which it has the highest *overlap score*. It is possible that a read can get aligned to multiple transcripts with the same *overlap score*. In such cases, the read is assigned to the transcript whose 3’ end is closest to the 3’end of the read. In cases where the 3’ end of the candidate transcripts are located in the same genomic position, the read is assigned randomly to one of them. Finally, the transcript is quantified by simply counting the number of reads assigned to it and converting these counts values to *Transcript Per Million* (TPM).

### Evaluation of quantification

We tested the quantification algorithm on simulated as well as real datasets. Throughout the paper, we use the widely accepted *Transcripts Per Million* as transcript and gene level metric for evaluation of predicted expression values. In addition, we also use the Mean Absolute Relative Difference (MARD) metric which was previously used for transcriptome comparisons [10], which we explain briefly below. For each transcript *i*, we calculate the *Absolute Relative Difference*:

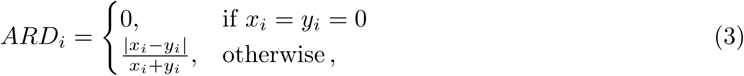

where *x*_*i*_ and *y*_*i*_ are the true value and estimated value respectively for transcript *i*. For the simulated dataset, the actual number of reads originating from a transcript *i* was considered as the true value and the number of reads predicted to have been originated from transcripts *i* by AERON was considered as the estimated value. However, in case of real datasets, the true count of the reads originating from a transcript is unknown. Hence, for read datasets, TPM value obtained using short reads for a transcript *i* was considered as the true value of *i* and the TPM value obtained using AERON was considered as the estimated value of *i*.

For a predicted value closer to the true value, the ARD values tends to be closer to 0. The *Mean Absolute Relative Difference* (MARD) for *M* transcripts is calculated simply by:

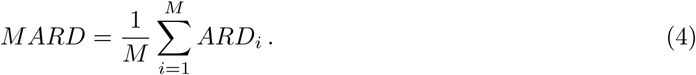

For comparison of expression estimates from long read datasets against the expression estimates obtained from short read data, we calculated the TPM values for all the transcripts present in the annotation file. Further, we obtained the gene-level quantification of a gene *g* by summing up the TPM values of all the transcripts belonging to *g*. We then calculated the Spearman correlation between the TPM values obtained from the long read dataset and the TPM values obtained from the short read data at transcript level. The Spearman correlation was also calculated at gene-level taking all the ENSEMBL(v92) annotated protein coding and non-coding genes into consideration.

### Fusion detection

Figure 4 shows an overview of the fusion detection pipeline, which we explain in the following.

#### Partial alignments

The reads are first aligned to the gene-exon graphs. This proceeds the same way as in the quantification pipeline, except that secondary alignments are not removed and the same part of a read may be mapped to multiple gene-exon graphs.

#### Tentative fusions

The partial alignments are used to create a list of *tentative fusions*. Whenever a read has a pair of partial alignments to two different genes whose endpoints are within 20 base pairs to each others in the read, the read supports a tentative fusion between the two genes. Each read may support multiple tentative fusions.

#### Fusion graphs

Each tentative fusion induces a *fusion graph*. The fusion graph combines the gene-exon graphs of the two participating genes. An extra crossover node is added to connect the two gene-exon graph. Each base pair in the first gene-exon graph is connected to the crossover node, and the crossover node is connected to each base pair in the second gene-exon graph. This way, the alignment may cross from any point in the first gene-exon graph to any point in the second gene-exon graph.

#### End-to-end fusion alignments

The reads are aligned to their fusion graphs. However, here the alignment must span the entire read from start to end. Clipped read ends are considered indels and contribute to the number of mismatches in the alignment.

#### End-to-end nonfusion alignments

Each read supported a list of tentative fusions previously. We extract the list of genes involved in those fusions for each read. In addition to this, we extract each pair of tentative fusions which include those genes regardless of which read supports the fusion, and say that these genes are relevant for the read. That is, if read *R*_1_ supports a fusion between genes *G*_1_ and *G*_2_ and nothing else, but an another read supports a fusion between *G*_2_ and *G*_3_, then all of *G*_1_, *G*_2_ and *G*_3_ are relevant for *R*_1_. The reads are then aligned to all of their relevant gene-exon graphs. Again the alignments must span the entire read from start to end.

#### Fusion scores

Once the reads have been aligned end-to-end to both the fusion graphs and the gene-exon graphs, the alignment scores are compared to calculate a *fusion score*. Given the lowest alignment edit distance to a fusion graph *C*_*f*_ and the lowest alignment edit distance to a gene-exon graph *C*_*n*_, the fusion score of a read to the fusion graph is defined as *C*_*n*_ − *C*_*f*_. This essentially describes how much better the read aligns to a fusion than any individual gene; a fusion score of 0 means that the read aligns to a fusion graph just as well as to a non-fusion graph, and higher fusion scores mean that the read aligns better to the fusion graph than any non-fusion graph. Note that the alignment edit distance to a fusion graph cannot be worse than the edit distance to a gene-exon graph, since the fusion graphs include the gene-exon graphs as subgraphs. The fusion graph alignment with the lowest edit distance is kept and the others are discarded. At this point each read can only support one fusion, which removes a large number of false positives.

#### Predicted fusions

The reads are filtered based on the error rate of the alignment to the fusion graph and the fusion score. Reads whose fusion score is below a user-given threshold (default 200) are discarded. Reads whose alignment error rate to the fusion graph is above 20% are also discarded. The paths of the fusion alignments are taken as the predicted fusion transcripts. When multiple reads align to the same fusion graph and cross over at the same exon, they are considered the same fusion event and one of them is arbitrarily selected as the fusion transcript. If multiple reads align to the same fusion graph but cross over at different exons, they are considered separate events. The output of this step is a list of predicted fusion transcripts.

#### Fusion support and alignments

Finally, all reads are aligned to the predicted fusion transcripts and the reference transcriptome using Minimap2 [19]. A read is then considered to support a fusion if its primary alignment crosses the fusion breakpoint with at least 150 base pairs on both sides. This recovers some reads which were missed by the earlier steps and removes some spurious fusions. The output of this step is a BAM file containing the alignments of the reads to the transcriptome and predicted fusion transcripts, and the number of reads that support each predicted fusion transcript.

The simulated data experiment produced 20 false positive calls even after the fusion score filtering. We recommend manually inspecting the predictions to filter out more false positives. In the K562 and NA12878 experiments, we curated the predictions by visualizing them with IGV [39] to remove cases where reads align poorly to one side of the fusion, and by aligning the predicted fusion transcripts to the reference genome and transcriptome using BLAST [48] to remove false positives caused by sequence similarity between transcripts.

### Simulation of ONT and parameter optimization

Two important parameters for AERON are the *seed length* and the *number of seeds* (NOS) for alignment of reads to the graph. Seed hits are exact matches between a part of the read and a part of a node, and are used for starting the alignments. Seed length is the minimum length of the exact matches. The number of seeds (NOS) denotes how many of the available seeds are used to compute an alignment between the read and the graphs. We wanted that the default values of these parameters in AERON should give us a good accuracy of the quantification in a reasonable runtime. For this, we first simulated 1M Oxford Nanopore reads using Nanosim on transcriptome mode (version 2.5.0, [49]). The novel K562 ONT data was given as the reference to Nanosim to create the training read profile. All the parameters of Nanosim was set to default. All the parameters of the algorithm were set as default. We performed several runs of AERON on the simulated dataset. Each run consisted of a different combination of seed length and the number of seeds. We measure the accuracy of quantification using the MARD score (See section —-). Figure 8 shows the effect of varying the seed length and the number of seeds on the runtime (*x*-axis) and the MARD score (*y*-axis). Each curve in the graph represents a single NOS parameter value and each point in the curve represents a seed-length parameter value. As expected, with the increase in the number of seeds accuracy of the quantication improves, evident by the decrease in the MARD score. Note, that MARD score measures the distance between the estimated value and the true value. Hence, lower the score, better the accuracy of the estimation. But setting the NOS parameter too high results in a higher runtime. We also observe that with the increase in seed length, the accuracy of the quantification goes down. For AERON, we selected a combination of parameters (NOS=15 and Seed-length=17) which was good trade-off between the accuracy and the runtime. Although, there might be other combination which might give similar or better results based on the dataset used in the pipeline.

**Figure 8.**
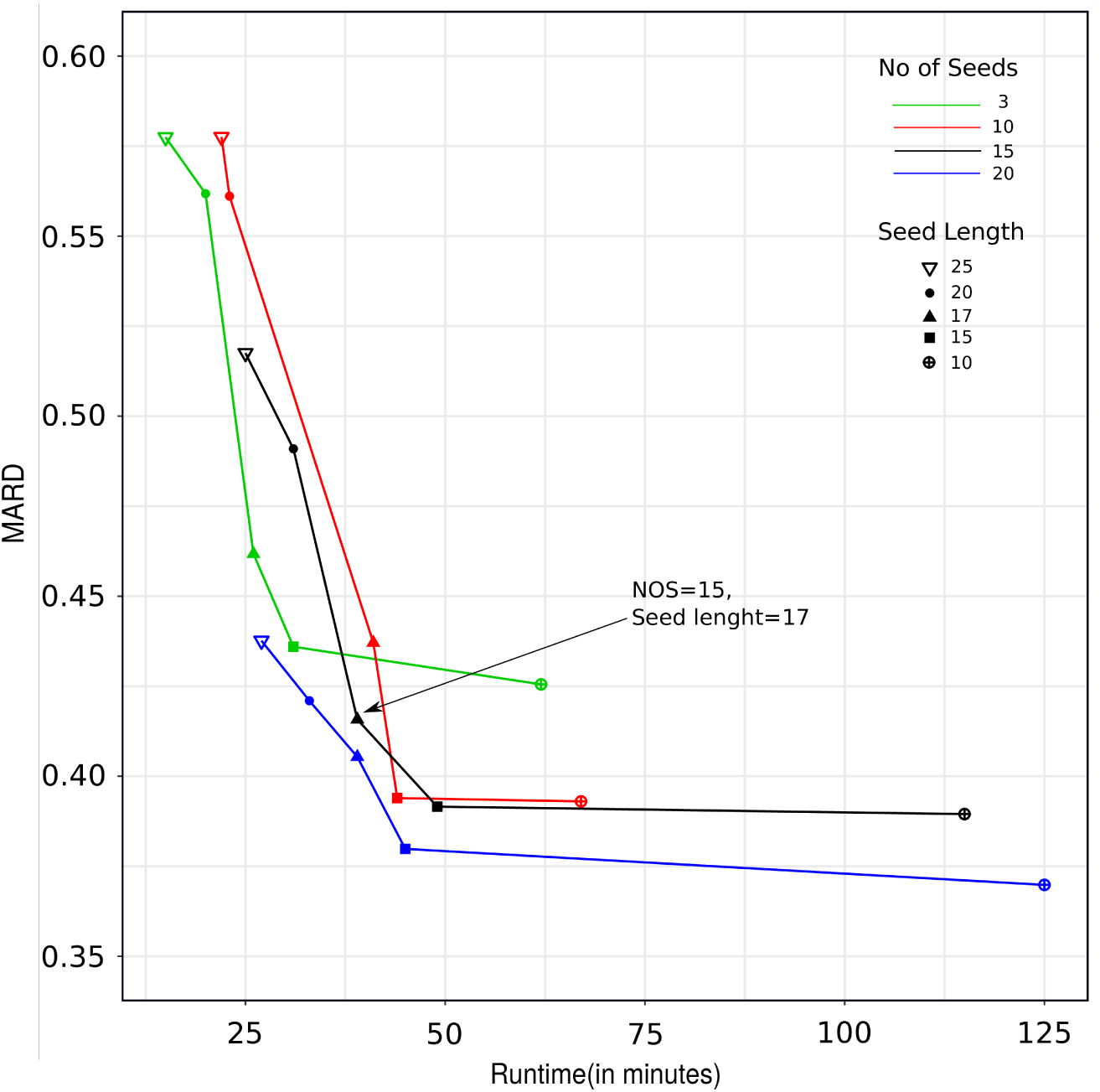
Comparison of the MARD score (y-axis) against runtime in minutes (x-axis) for different parameter configurations of AERON on simulated data. Each line in the plot represents a value for the *No. of Seeds* parameter. Each point within a line represents a value of the*Seed Length* parameter. One such value is marked with an arrow, which denotes the default value of Aeron.

### Datasets

Two different Oxford Nanopore Technology (ONT) read sets were used for the analysis: a novel dataset with 2.7M reads from K562 cells with an average read length of 750bps, and an available dataset with 25M reads from NA12878 cells with average read length of 1030bps [50]. For the alignment step of AERON, human genome from GRCh38.p12 [51] was used as the reference. We also aligned the two datasets against the human reference genome using Minimap2 and filtered out all the secondary alignments keeping only the primary alignments. For each transcript in the genome, we then counted the number of reads aligned to the transcript and converted the numbers into TPM values.

To compare the expression estimates from AERON and Minimap2 against expression estimates from short reads, we downloaded two short read data-sets: 113M paired end reads from the K562 cell line (SRX4958124, [52]) and 188M paired end reads from NA12878 (ERX329208, [53]). We then calculated the expression for the two datasets using Salmon ([10], v0.11.2) with default parameters except *k*mer size, which was set to 17.

## Availability of data and materials

Data used for experiments can be accessed via the accession numbers given in the Methods section of the manuscript. The algorithm is freely available under MIT license and can be accessed via https://github.com/SchulzLab/Aeron

## Author’s contributions

MR, DD, JG, MS and TM conceived the idea. DD implemented the gene-exon graph construction and performed the quantification analysis. MR implemented the graph alignment and the fusion pipeline, and performed the fusion analysis. DD and MR implemented the expression quantification pipeline. CY performed the 3’ distance analysis. LX and HML generated the K562 data. MR, DD, JG, MS and TM wrote the manuscript with input from all authors.

## Acknowledgments

This work has been supported by the DZHK (German Centre for Cardiovascular Research, 81Z0200101) and the DFG Clusters of Excellence on Multimodal Computing and Interaction [EXC248] and Cardio-Pulmonary Institute (CPI) [EXC 2026].

