## Supplementary for "AERON: Transcript quantification and gene-fusion detection using long reads"

Rautiainen *et al*

Abstract

Single-molecule sequencing technologies have the potential to improve measurement and analysis of long RNA molecules expressed in cells. However, analysis of error-prone long RNA reads is a current challenge. We present AERON for the estimation of transcript expression and prediction of gene-fusion events. AERON uses an efficient read-to-graph alignment algorithm to obtain accurate estimates for noisy reads. We demonstrate AERON to yield accurate expression estimate on simulated and real datasets. It is the first method to reliably call gene-fusion events from long RNA reads. Sequencing the K562 transcriptome, we used AERON and found known as well as novel gene-fusion events.

### 1 Graph construction

Below we summarize the graph construction procedure for AERON (Algorithm **??**). Given a gene $g$ and the set of all transcripts belonging to $g$, we begin by generating the border list $\rho(g)$ and its sub lists namely acceptor list $\alpha(g)$ and donor list $\delta(g)$ (Line 3). We also create a list of the position of all the exonic bases present in $g$ and name it as *exon-list* $\chi(g)$ (Line 4). This provides us with all the splice sites present in the genome. Each exon of the gene forms a vertex in the graph and each vertex is split if the exon corresponding to the vertex has an alternative splice site. This is implemented in the following steps. We create a string $s$ and initialize it with the first position listed in $\alpha$ ($\alpha[0]$) of $g$ (Line 6). Starting from the next position $i = \alpha[0] + 1$, we check whether the base at position $i$ is an *exonic-base.* If yes then we concatenate the base to $s$ (Line 10-11). We continue this procedure till we encounter an $i$, which is a member of either a 5'-border or a 3' border (Line 12 and Line 17). Encountering a 5'-border indicates an alternate acceptor site present in the exon. Hence, in concurrence to equation (**??**), we consider the string $s$ in its current form as a vertex and add it to $V$. We label the vertex with an integer $i - 1$. To keep a track of order in which the vertices were created we add the label to a list $L$ (Line 14). We will use this information later for adding edges between the vertexes. We reset $s$ to the base at the $i^{th}$ position (lines 12-16). A 3' border indicates the end of an exon. Hence, when a 3' border is encountered, we concatenate the base at the $i^{th}$ position to $s$ and consider it as a vertex with label $i$. We add the vertex to $V$ and add the label to list $L$. We reset the string $s$ to an empty string (lines 17-21). While traversing, if we encounter a non-exonic base, we skip the position and jump to the next 5' border in the gene sequence which has not been encountered yet and continue our iteration (lines 22-25). The above steps are repeated till all the borders in $g$ are encountered. We then traverse the vertex set $V$. Given two vertices $v_i$ and $v_j$ belonging to $V$ where $i$ and $j$ are vertex labels, we add an edge between $v_i$ and $v_j$ if $i < j$ (lines 28-33).

This procedure is repeated for all the genes in the reference species.

**Algorithm 1** Graph construction for AERON

**Input:** gene sequence $g$, transcripts set $\mathcal{T}_g$.
S = *obtainSites(g)*
*generateList*(S, $\rho(g)$, $\alpha(g)$, $\delta(g)$)
*generateList*($\chi(g)$)
$V = E = \phi$
$s = g[\delta[0]]$
$c = 1$
$L = []$
**for** $i \in (\alpha[0] + 1, \delta[-1])$ **do**
    **if** $i \in \chi$ **then**
        $s = s + g[i]$
    **else if** $i \in \delta$ **then**
        $addNode(s, V, i - 1)$ → Adding node sequence $s$ with label $i - 1$ to $V$
        *append(L,i-1)*
        $s = g[i]$
        $c = c + 1$
    **else if** $i \in \rho$ **then**
        $s = s + g[i]$
        $addNode(s, V, i)$ → Adding node sequence $s$ with label $i$ to $V$
        *append(L, i)*
        $s = NULL$
    **else**
        $i = \delta[c]$
        $s = g[\delta[c]]$
        $c = c + 1$
    **end if**
**end for**
**for** $i \leftarrow 1, |L|$ **do**
    $n_i$ = *getNode*$(V, L[i])$ → obtaining the node having label $i$
    **for** $j \leftarrow i + 1, |L|$ **do**
        $n_j$ = *getNode*$(V, L[j])$ → obtaining the node having label $j$
        $E$ = *addEdge*$(n_i,\ n_j)$
    **end for**
**end for**
**Output:** $G_g$=$(V, E)$

#### 2 Alignment of reads along the transcripts

Recent studies have shown that during Oxford Nanopre sequencing, most of the reads originate near the 3' end of a transcript [?]. To test this in our datasets we aligned the reads to the combined annotations of coding and non-coding RNAs (Hg38 Ensembl version 91) using minimap2 version 2.17 [?]. The parameters were set as "-ax map-ont -t 3". Addtitonally, we set "-uf" parameter for direct RNA sequencing runs. We calculated the distributions of read distances to 5' and 3' ends of isoforms using the primary alignments with read distances adjusted by the lengths of transcripts. As expected, we found that most of the reads align near the 3' end of the transcript(**??** and **??**). The behaviour remains constant irrespective of the sequencing protocol (Fig.**??**).

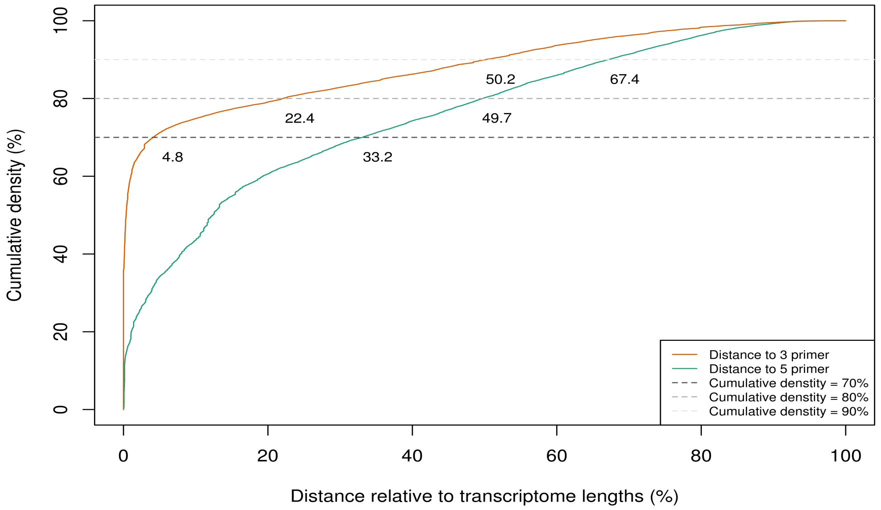


Figure S1: Cumulative density distribution of reads along the length of the transcripts for K562. Here, we observe an alignment bias towards the 3' end of the transcripts. In other words, most of the reads are generated near the 3' end of a transcript.

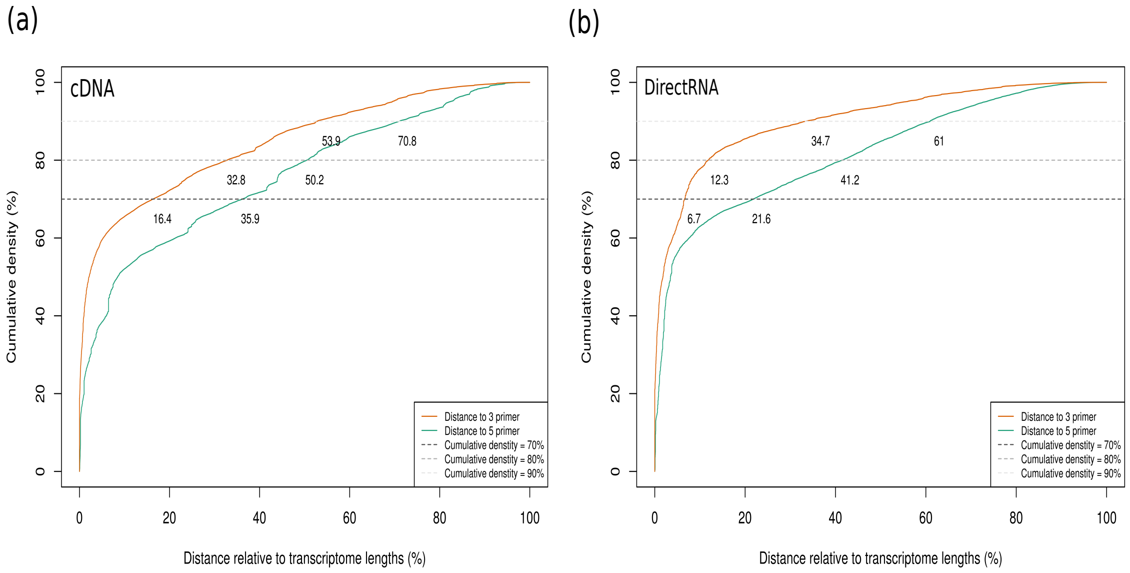


Figure S2: Cumulative density distribution of reads along the length of the transcripts for NA12878 data sequenced from (a) complementary DNA and (b) directly from native RNA. Like in K562 dataset, we observe an alignment bias towards the 3' end of the transcripts.

### 3 Subset analysis

To check whether AERON produces reproducible quantifications, we divided 25M NA12878 reads into three subsets consisting of 8M reads in two subsets and 9M reads in the third subset. We executed the quantification step of AERON on all the three subsets seperately with default parameters and calculated the gene level and transcript level expression estimates. We found that both the gene level and transcript level estimates are highly correlated across (Fig. ?? and ??) subsets with the spearman correlation of approximately 0.97 at gene level and a correlation of 0.92 at transcript level. This implied that AERON was able to produce reproducible expression across various sub-samples.

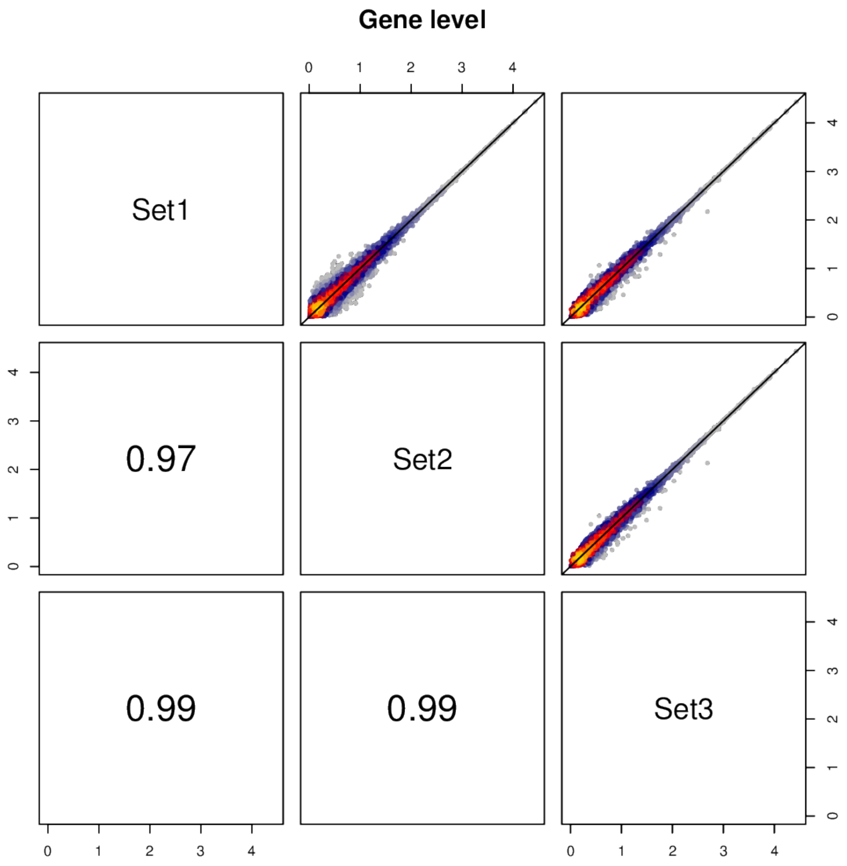


Figure S3: Gene level quantification comparison across three random subsets of NA12878 datasets.

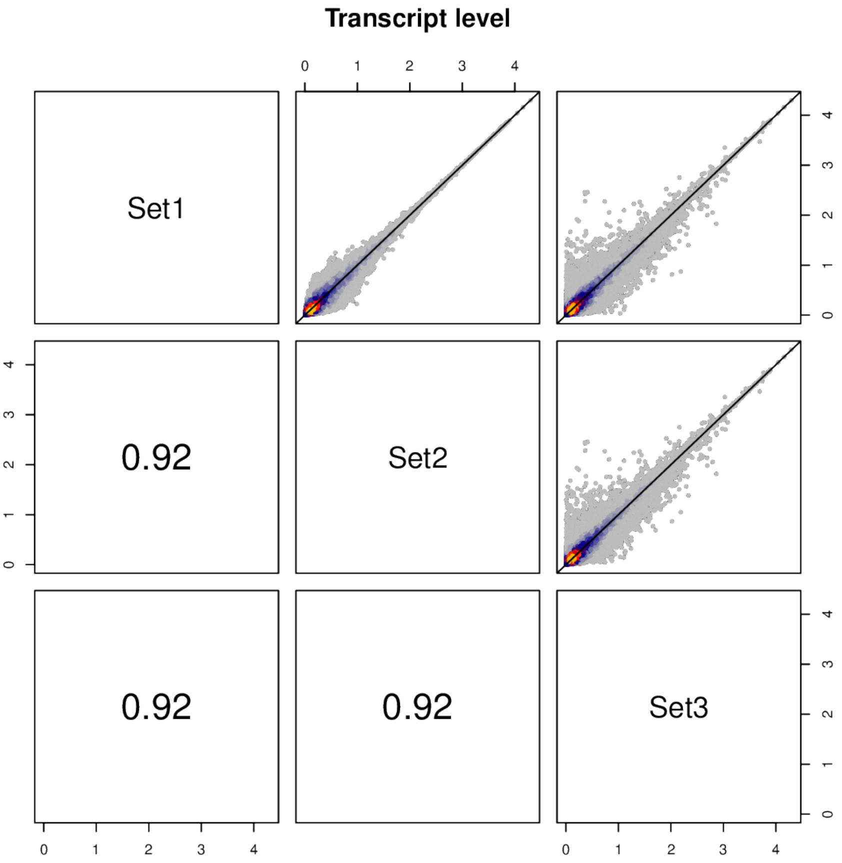


Figure S4: Transcript level quantification comparison across three random subsets of NA12878 datasets.

#### 4 Average base quality and average read length of K562 and NA12878 datasets

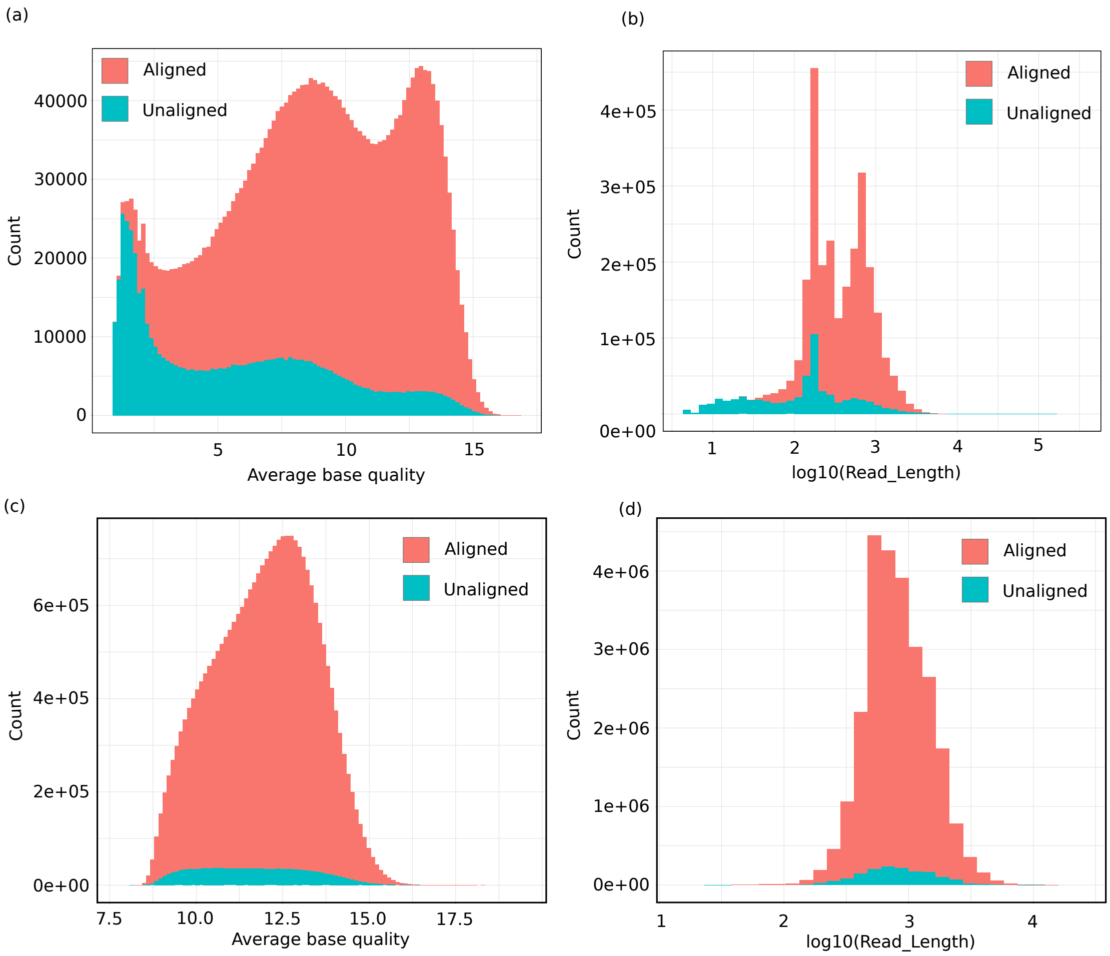


Figure S5: Base quality distribution(a and c) and read length distribution (b and d) of aligned and unaligned reads from K562 (top row) and NA12878 (bottom row) datasets. We observe here that most of the unaligned reads are short in length and consists of a large number of low quality bases.

| Dataset | AERON | | | Minimap | | |
|---|---|---|---|---|---|---|
| | #reads mapped | Correlation | MARD | #reads mapped | Correlation | MARD |
| K562 (2.7M) | 2167286 | **0.297** | **0.635** | 1074411 | 0.209 | 0.659 |
| NA12878 (25M) | 23902112 | **0.270** | 0.664 | 23211716 | 0.207 | **0.637** |

Table S1: Spearman correlation and MARD between Transcripts Per Million (TPM) at transcript level obtained from AERON/Minimap using Oxford Nanopore Sequencing (ONT) data and TPM at transcript level obtained from Salmon using Illumina data

### 5 Quantification of K562 and NA12878 datasets at transcript level

We found that the correlation between the expression estimates from AERON and expression estimates using short reads (obtained using salmon [?]) was consistently low (Tab.**??**. Presence of small length transcripts could be one possible reason for this behavior. The error prone reads generating from these small length transcripts might not get aligned correctly and hence the transcript might not get quantified. To test this hypothesis, we removed transcript shorter than a defined cut-off and then calculated the spearman correlation using the rest of the transcripts. As expected, with the increase in the cut-off length, the correlation between the expression estimates from AERON and the expression estimates from short reads improved significantly (Fig.**??**).

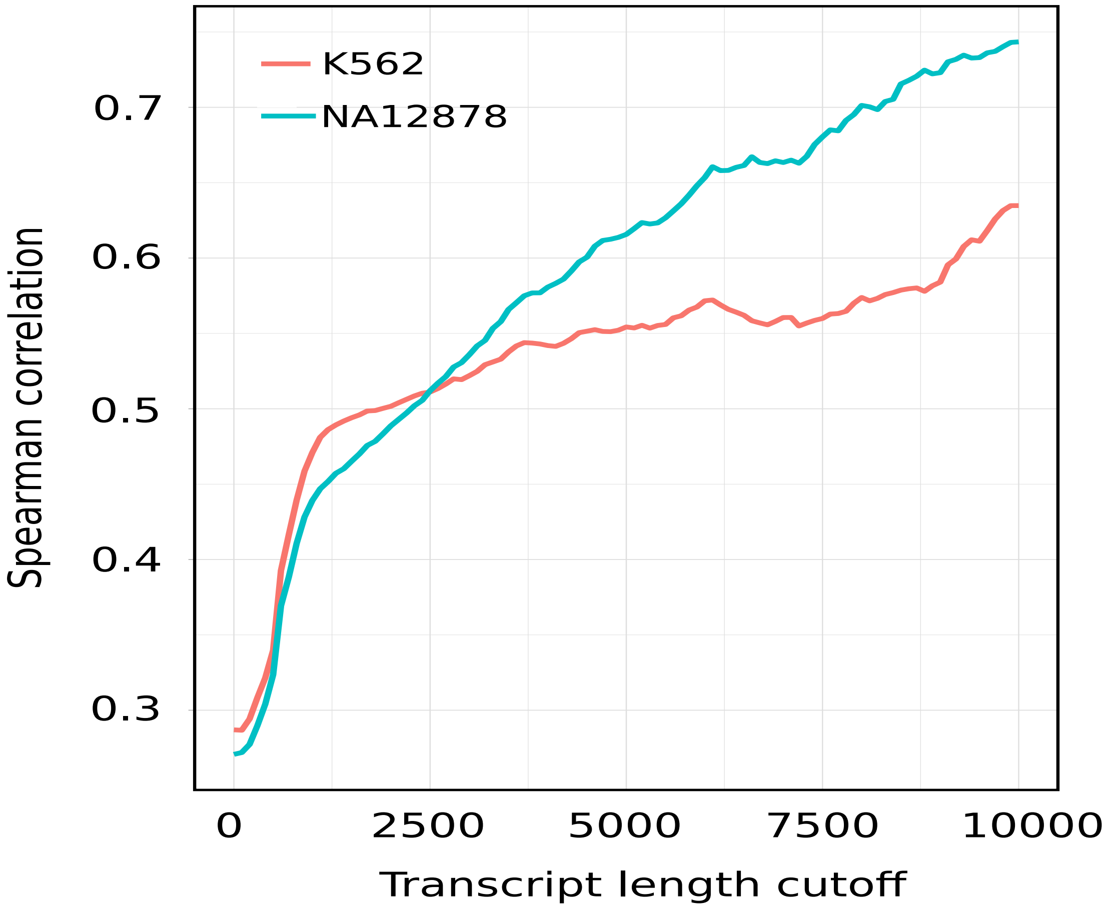


Figure S6: Effect of transcript length cutoff on spearman correlation between the expression estimates from AERON and expression estimates from salmon

### 6 Fusion events predicted by AERON

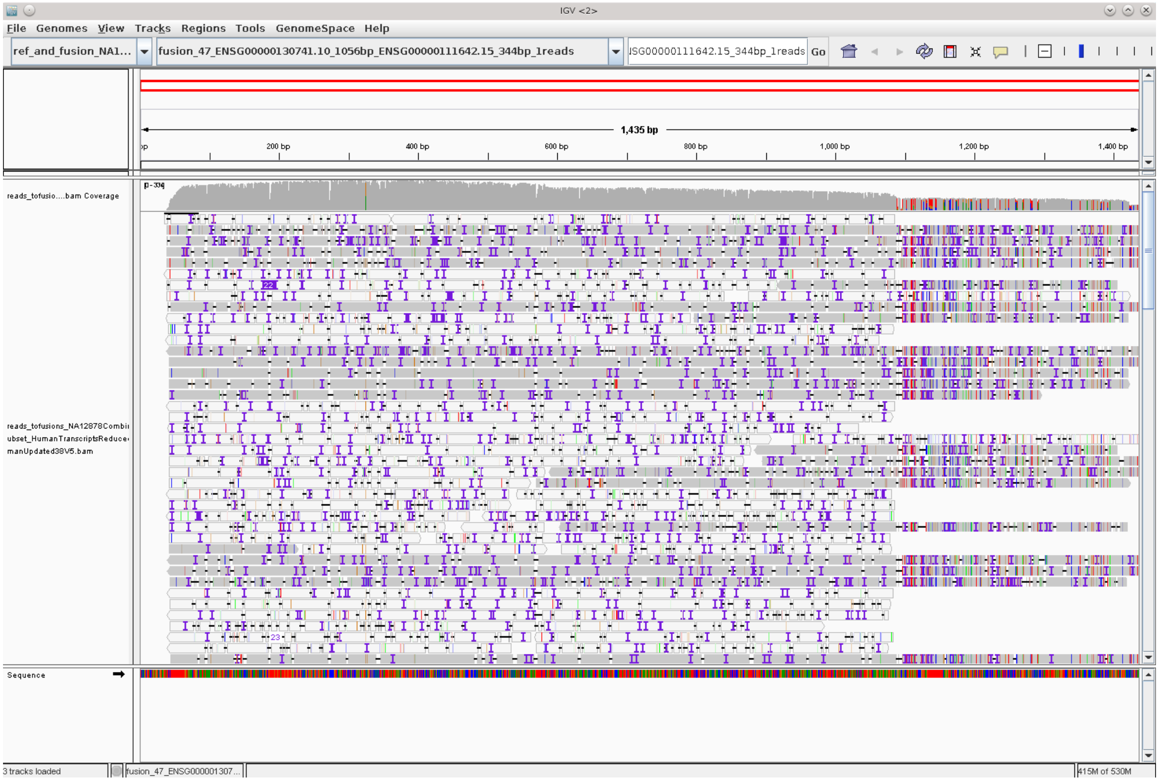


Figure S7: An IGV screenshot of a predicted fusion transcript that was rejected by manual curation. The fusion breakpoint is around 1100 bp. The alignments on the right side are clearly worse than the alignments on the left side, showing that the fusion is likely to be a false positive.

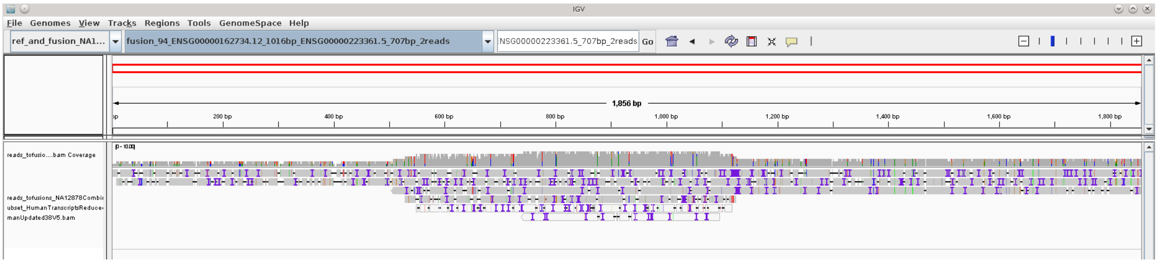


Figure S8: An IGV screenshot of the predicted FTH1P10-PEA15 fusion transcript from NA12878. The fusion breakpoint is around 1130 bp.

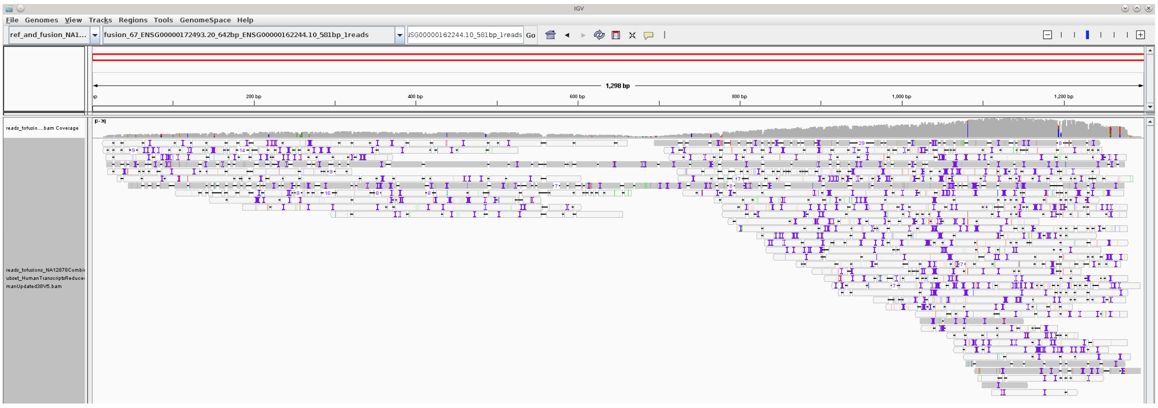

Figure S9: An IGV screenshot of the predicted AFF1-RPL29 fusion transcript from NA12878. The fusion breakpoint is around 700 bp.

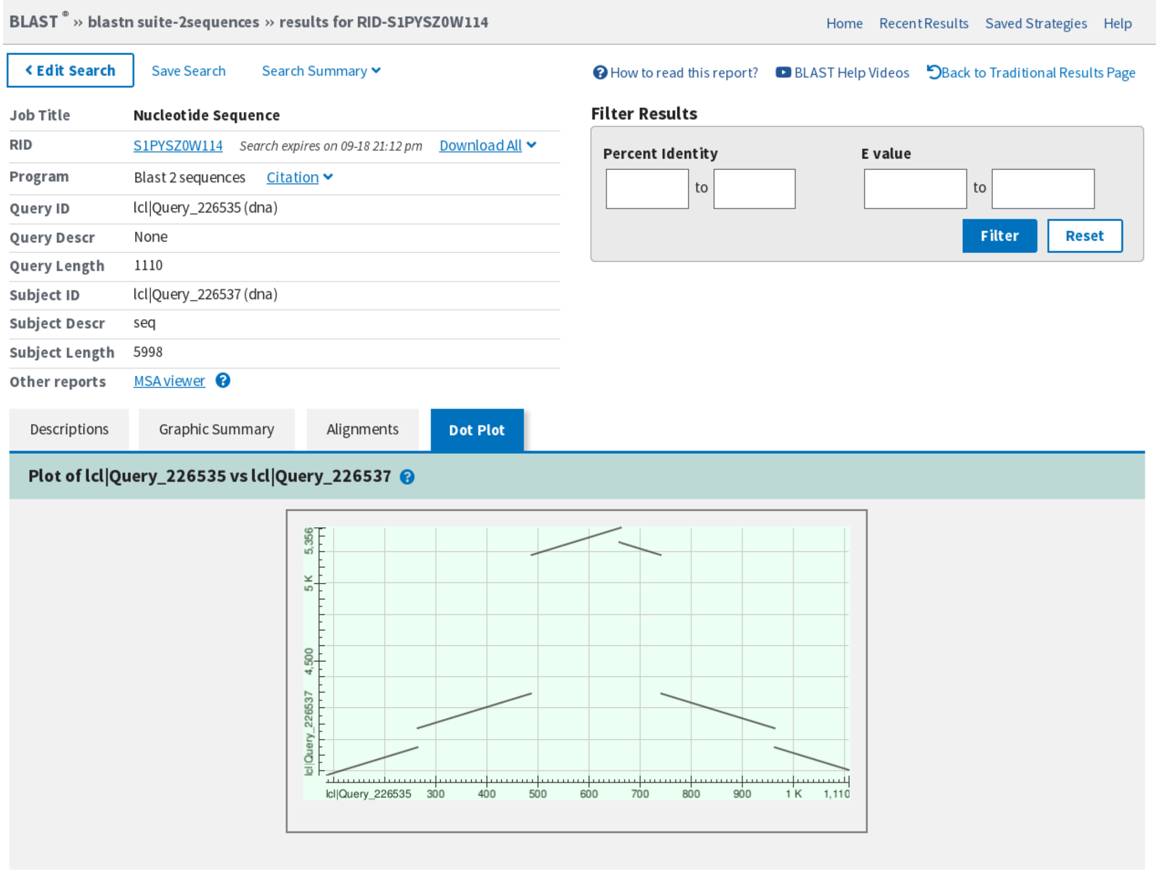


Figure S10: A BLAST screenshot showing the dot plot of the alignment between the predicted HBG2-HBG1 fusion event in the K562 data (x-axis) and chromosome 11 around 5,499,607-5,505,605 (y-axis) which contains the end of the HBG2 gene. The alignment is consistent with an inverted duplication of the region.

| Dataset | Aeron | | Minimap2 | |
|---|---|---|---|---|
| | Memory(in GB) | Runtime(HH:MM:SS) | Memory(in GB) | Runtime(HH:MM:SS) |
| K562 (2.7M) | **4.97** | 01:02:21 | 5.64 | **00:17:29** |
| NA12878 (25M) | **5.27** | 11:15:25 | 7.12 | **04:29:10** |

Table S2: Runtime and memory comparison between the runs of AERON and Minimap

Transcript Level

| Dataset | Aeron | Minimap2 | Salmon (Quasi-Mapping) | Salmon (Alignment-based) |
|---|---|---|---|---|
| K562 (2.7M) | **0.297** | 0.209 | 0.233 | 0.215 |
| NA12878 (25M) | **0.276** | 0.233 | 0.257 | 0.221 |
| Gene Level | | | | |
| K562 (2.7M) | **0.836** | 0.704 | 0.706 | 0.700 |
| NA12878 (25M) | **0.823** | 0.778 | 0.759 | 0.772 |

Table S3: Alternate approaches for quantifying transcripts using long reads. The table shows the spearman correlation between the expression estimates obtained using short reads (using salon) against expression estimates obtained using AERON(column 2), only minimap2 output(column 3), salmon in quasi mapping mode with long reads as input (column 4) and salmon in alignment-based mode on output generated from minimap2 (column 5). The numbers of reads

### 7 Alternate approaches for quantifying transcripts using long reads

Soneson *et al.* [?] recently proposed an alternate approach for quantifying transcripts using the EM based approach Salmon[?] in two modes. In the first mode, they ran salmon in quasi-mapping settings with the index generated from ENSEMBL cDNA reference fasta file and in the second mode they ran salmon in alignment-based settings on the output bam file produced by minimap. We tested the suggested approaches on out datasets and found no major effect on the transcript level expression estimates.
